# DNA demethylation is a driver for chick retina regeneration

**DOI:** 10.1101/804161

**Authors:** Agustín Luz-Madrigal, Erika Grajales-Esquivel, Jared Tangeman, Sarah Kosse, Lin Liu, Kai Wang, Andrew Fausey, Chun Liang, Panagiotis A. Tsonis, Katia Del Rio-Tsonis

**Affiliations:** Department of Biology and Center for Visual Sciences at Miami University, Miami University, Oxford, OH 45056, USA.; Department of Biology and Center for Stem Cell & Organoid Medicine (CuSTOM), Cincinnati Children’s Hospital Medical Center, Cincinnati, OH 45229, USA; Department of Computer Science and Software Engineering, Miami University, Oxford, OH 45056, USA; Department of Biology, University of Dayton and Center for Tissue Regeneration and Engineering at the University of Dayton (TREND), Dayton, OH 45469, USA.

## Abstract

**Background:** A promising avenue toward human retina regeneration lies in identifying the factors that promote cellular reprogramming to retinal neurons in organisms able to undergo retina regeneration. The embryonic chick can regenerate a complete neural retina, after retinectomy, via retinal pigment epithelium (RPE) reprogramming in the presence of FGF2. Cellular reprogramming resets the epigenetic landscape to drive shifts in transcriptional programs and cell identity. Here, we systematically analyzed the reprogramming competent chick RPE prior to injury, and during different stages of reprogramming. We examined the dynamic changes in the levels and distribution of histone marks and DNA modifications, as well as conducted a comprehensive analysis of the DNA methylome during this process.

**Results:** In addition to changes in the expression of genes associated with epigenetic modifications during RPE reprogramming, we observed dynamic changes in histone marks and intermediates of the process of DNA demethylation. At early times after injury, H3K27me3 and 5mC repression marks decreased while 5caC and the H3K4me3 activation mark increased, suggesting genome-wide changes in the bivalent chromatin, impaired DNA methylation, and active DNA demethylation in the chromatin reconfiguration of reprogramming RPE. Comprehensive analysis of the methylome by whole-genome bisulfite sequencing (WGBS) confirmed extensive rearrangements of DNA methylation patterns including differentially methylated regions (DMRs) found at promoters of genes associated with chromatin organization and fibroblast growth factor production. In contrast, genes associated with early RPE reprogramming are hypomethylated in the intact RPE and remain hypomethylated during the process. During the generation of a neuroepithelium (NE) at later stages of reprogramming, decreased levels of H3K27me3, 5mC, and 5hmC coincide with elevated levels of H3K27Ac and 5caC, indicating an active demethylation process and genome-wide changes in the active regulatory landscape. Finally, we identify Tet methylcytosine dioxygenase 3 (TET3) as an important factor for DNA demethylation and retina regeneration in the embryonic chick, capable of reprogramming RPE in the absence of exogenous FGF2.

**Conclusion:** Our results demonstrated that injury signals early in RPE reprogramming trigger genome-wide dynamic changes in chromatin, including bivalent chromatin and DNA methylation. In the presence of FGF2 these dynamic modifications are further sustained in the commitment to form a new retina. We identify DNA demethylation as a key process driving the process of RPE reprogramming and identified TET3 as a factor able to reprogram RPE in absence of FGF2. Our findings reveal active DNA demethylation as an important process that may be applied to remove the epigenetic barriers in order to regenerate retina in mammals.

## Background

Stem cell therapy and induced retina regeneration constitute potential strategies to reach the ultimate goal of regenerative medicine to repair lost or damaged retina [1, 2]. Age-related macular degeneration (AMD) and diabetic retinopathy are common among the visually impaired population [3]. After damage or disease, humans do not possess the intrinsic capability to regenerate their retina. In contrast, several non-mammalian vertebrates, including the chick, have the potential to regenerate their entire retina from endogenous cell populations after retinal damage [4, 5]. The embryonic chick can regenerate its retina at embryonic day (E) 4-4.5 (stages 23-25; [6, 7]) by reprogramming cells of the retinal pigment epithelium (RPE) into retina progenitor cells that eventually differentiate into the major retinal cell types [8–11]. We previously demonstrated that during the Phase I of RPE reprogramming, within 6 hours after surgical removal of the retina (retinectomy), the quiescent cells of the RPE transiently reprogram (these cells are referred to as transient reprogrammed RPE, t-rRPE), and express pluripotency inducing factors including *sox2*, *c-myc*, *klf4*, and eye field transcription factors (EFTF), while simultaneously down-regulating RPE specific markers [11]. In the same study, we showed that FGF2 is necessary to complete the RPE reprogramming process (cells referred to as reprogrammed RPE, rRPE), sustaining the expression of EFTF and pluripotency inducing factors, as well as upregulating the expression of *lin-28a* in the RPE. Later during the Phase II, the rRPE proliferates to generate a NE containing retina progenitor cells [11]. RPE reprogramming is not spontaneous after injury, as it requires an exogenous source of FGF2. Therefore, chick retina regeneration represents a powerful model to study individual steps associated with reprogramming including injury (t-rRPE), dedifferentiation, and differentiation in the presence of FGF2 (rRPE).

During cell reprogramming, including somatic cell nuclear transfer, cell fusion, and generation of induced pluripotent stem cells (iPSC), the epigenetic landscape is reset, adopting a conformation of open chromatin. This “flexible” chromatin state allows transcription programs and cell identity to shift [12, 13]. Chromatin state can be influenced by DNA methylation, hydroxymethylation, and histone modifications at specific genomic regions, and has an important impact on epigenetic regulation of gene expression and cell specification during development and reprogramming [14–18]. DNA methylation can regulate gene expression by maintaining the silent state of chromatin in time and tissue-specific manners [18]. Generation of the methylated form of cytosine, 5-methylcytosine (5mC), is catalyzed by DNA methyltransferases (DNMTs). DNMT3a and DNMT3b are involved in *de novo* methylation, whereas DNMT1 is responsible for maintaining DNA methylation patterns during DNA replication [18]. In contrast, active DNA demethylation is a multistep process mediated by ten-eleven translocation dioxygenases (TET) that can sequentially oxidize 5mC to 5-hydroxymethylcytosine (5hmC), 5-formylcytosine (5fC), and 5-carboxylcytosine (5caC). Thereafter, 5fC or 5caC can be excised by thymine DNA glycosylase (TDG) and the resulting apyrimidinic sites can be replaced by unmodified cytosine through base excision repair (BER) [17].

Histone modifications constitute another regulatory mechanism involved in the control of gene expression. In human and mouse embryonic stem (ES) cells, large regions of repressive mark trimethylated histone H3 lysine 27 (H3K27me3) co-exist with smaller regions of active mark trimethylated histone H3 lysine 4 (H3K4me3) [19–22]. These regions, called bivalent chromatin, are present in promoters of transcription factors with importance in development and differentiation [19, 20]. During differentiation, bivalent promoters tend to preserve the activation or repression mark, but not both, suggesting that bivalent domains are kept in a poised state in ES cells [19]. Trithorax (TrxG) and Polycomb group (PcG) proteins are the key players in maintaining bivalent domains [23]. PcG proteins form two multi-subunit repressive complexes called Polycomb Repressive Complexes 1 and 2 (PRC1/2) that are involved in gene silencing of many developmental processes [24, 25]. Histone 3 lysine 27 di-/tri-methylation (H3K27me2/3) is catalyzed by the enzymatic subunit SET domain-containing protein Enhancer of zeste homologue 2 (EZH2) present in PRC2 [24]. Contrary to PcG, TrxG maintains transcriptionally active chromatin and forms a complex in which the methylase Mll1 catalyzes H3K4me3 modifications [26]. H3K27 methylation is considered to be relatively stable and maintains long-term transcriptional repression; however, lysine demethylases such as JMJD3 (Jomonji domain containing 3, Kdm6b) and UTX (Kdm6a) specifically demethylate H3K27, resulting in activation of genes associated with animal body patterning, inflammation, and ultimately resolution of bivalent domains [27].

Epigenetic mechanisms associated with regeneration have been partially investigated in organisms that have the capability to regenerate their tissues after injury [28–33]. For example, in the *Xenopus* froglet, DNA methylation affects the limb regenerative capacity mediated by an enhancer sequence named Mammalian Fish Conserved Sequence 1 (MFCS1) which controls the expression of *Shh* [34]. On the other hand, dynamic changes in DNA methylation and expression patterns of DNMTs have been observed during Müller glia (MG) reprogramming [35, 36] and in zebrafish fin regeneration [37–39]. It has also been demonstrated that MG reprogramming, proliferation, and optic nerve regeneration are affected after knockdown (KD) of apobec2a and apobec2b-enzymes involved in deamination of methylated DNA [40]. However, changes in DNA methylation during MG reprogramming are independent of Apobec2 expression [35], suggesting that other factors may play a role in DNA demethylation during MG reprogramming.

In addition to DNA methylation, histone modifications have been reported to be associated with the process of regeneration. In this regard, loss-of-function studies demonstrated that H3K27me3 demethylase KDM6B.1 is necessary for the process of fin regeneration in zebrafish [41]. Furthermore, genes encoding histone methyltransferases and acetyltransferases, in addition to histone modifications H3K9me2, H4K20me3, H3K4me3, and H3K14Ac, are differentially regulated or modified in this process [41, 42]. On the other hand, an Ezh2-deficient zebrafish line depicts defective spinal cord regeneration, suggesting that H3K27me3 and Ezh2 are important in this regenerative process [43]. In mammals, H3K27me3 demethylases JMJD and UTX are necessary for murine skin repair [44]. More recently it was suggested that DNA methylation and repressive chromatin may be the obstacles murine MG and RPE have to overcome for a regenerative program to ensue [45, 46].

While previous work suggests that chromatin modifications are associated with the process of regeneration, the mechanisms and key players of these processes are only partially known. In the present study, we performed a systematic analysis of chick RPE reprogramming process including differential gene expression of factors associated with epigenetic modifications, high-resolution and three-dimensional (HR-3D) reconstruction confocal microscopy of histone marks and DNA modifications, as well as whole-genome bisulfite sequencing (WGBS). Our results demonstrate differential regulation and genome-wide dynamic changes in DNA and histone modifications in addition to changes in the regulation of genes associated with epigenetic marks, more importantly DNA methylation and demethylation. During the Phase I of RPE reprogramming, at early times post-retinectomy (PR), H3K27me3 and 5mC repression marks decrease while H3K4me3 activation mark and intermediates of DNA demethylation 5caC and 5hmC increase, suggesting significant changes in bivalent chromatin, impaired DNA methylation, and active DNA demethylation in the t-rRPE in response to injury and in the rRPE committed to reprogramming in response to FGF2. Comprehensive analysis of the methylome by WGBS confirmed extensive re-arrangements of DNA methylation patterns in the t-rRPE and rRPE at early times PR. This was evident by differentially methylated regions (DMRs) found at the promoter of genes involved in regulation of chromatin organization and fibroblast growth factor production. In contrast, genes associated with reprogramming are hypomethylated in the intact RPE and remain hypomethylated during the process. Further, during the Phase II of RPE reprogramming, at 3 days PR (dPR), following the generation of a NE originated from the rRPE, decreased levels of H3K27me3, 5mC and 5hmC, coinciding with elevated levels of H3K27Ac and 5caC, suggest active demethylation and genome-wide changes in the active regulatory landscape during rRPE to NE. Finally, TET3 overexpression was sufficient to reprogram RPE in the absence of FGF2, demonstrating that the process of DNA demethylation and TET3 play a key role in RPE reprogramming. Our findings reveal a complex rearrangement in the levels and patterning of histones and DNA epigenetic marks associated with RPE reprogramming, and identify TET3 as a novel and sufficient factor to promote retina regeneration in the embryonic chick.

## RESULTS

### Histone and DNA modifications present in the reprogramming-competent RPE

To determine the presence and distribution of histone and DNA modifications at the time when the RPE is competent to undergo reprogramming (embryonic stages 23-25; [6]), we performed immunofluorescence staining of epigenetic marks associated with bivalent chromatin (H3K27me3 and H3K4me3), transcriptionally active chromatin (H3K4me2 and H3K27Ac), and DNA modifications, including 5mC and the oxidized forms 5hmC and 5caC. Repressive and active histone marks were widely distributed throughout the lens (L), pigmented epithelium (PE) and non-pigmented epithelium (NPE) of the ciliary margin (CM), optic cup lip (OCL), NE, and RPE (Figure 1a-d and h-k). The repressive mark H3K27me3 was less prevalent, with few scattered positive cells in the most posterior part of the NE and more enrichment in the presumptive retinal ganglion cells (P-RGCs), the first cell type to be specified during retinal neurogenesis [47] (Figure 1h). In contrast, the activation marks H3K4me3 and H3K4me2 were uniformly present in all different regions of the eye (Figure 1b, c and i, j). In the RPE, the activation mark H3K27Ac was more enriched in the most anterior part of the eye in comparison to the posterior region (Figure 1d, k). All DNA modifications, 5mC, 5hmC, and 5caC, were also present in the anterior and posterior regions of the eye (Figure 1e-g and l-n). Interestingly, similar to H3K27me3, 5hmC positive cells were less prominent in the most posterior region of the NE when compared to the anterior region of the NE (compare Figure 1f with 1m) and highly enriched in P-RGCs (Figure 1m), suggesting that H3K27me3 and 5hmC could be involved in the process of RGCs differentiation. Furthermore, a strong signal of 5hmC was observed in the RPE and CM including the OCL that contains multipotent optic cup stem/progenitor cells [48, 49] (Figure 1f, m). These results demonstrate the presence of bivalent histone marks and DNA modifications, including those associated with the process of demethylation (5hmC and 5caC) in the reprogramming-competent chick RPE.

**Figure 1.**
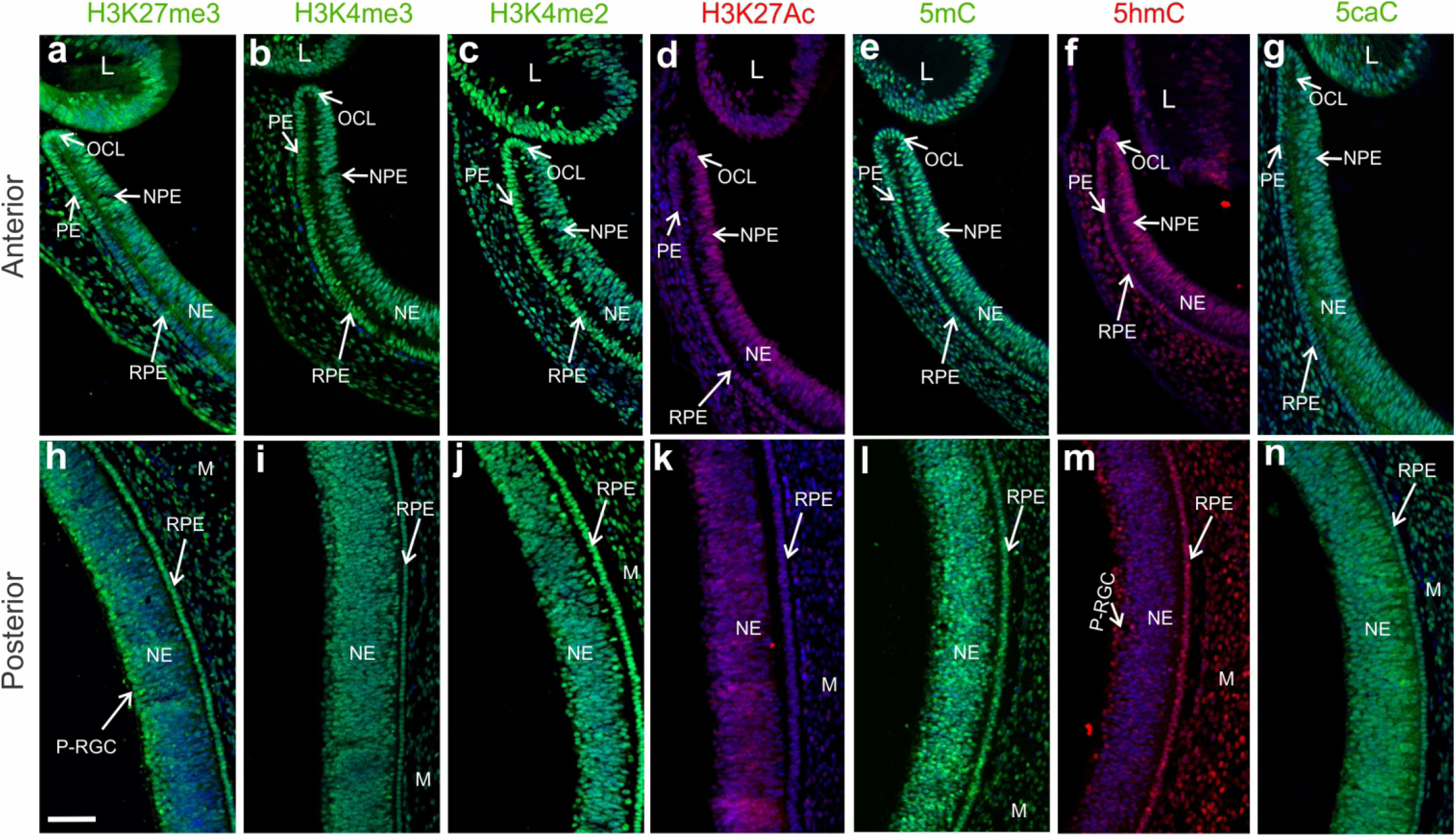
Histone and DNA modifications present in the reprogramming competent RPE. Immunofluorescence staining of histone modifications associated with bivalent chromatin (H3K27me3/H3K4me3) and activation marks H3K4me2 and H3K27Ac as well as DNA modifications 5mC, 5hmC and 5caC in the **(a-g)** anterior and **(h-n)** posterior regions of the chick eye at St 23-25 (E4). DAPI in blue, was used for nuclear counterstaining. NE: neuroepithelium; NPE: non-pigmented epithelium of the ciliary margin; PE: pigmented epithelium of the ciliary margin; RPE: retinal pigment epithelium; L: lens; M: mesenchyme; OCL: optic cup lip. P-RGC: Presumptive Retinal Ganglion Cells. Scale bar in **h** is 50 μm and applies to all the panels.

### Differential expression of genes associated with histone and DNA modifications in the t-rRPE and rRPE

In order to identify differentially expressed factors associated with histone marks and DNA modifications in t-rRPE and rRPE, we used laser capture microdissection (LCM) to collect intact RPE at stages 23-25, t-rRPE at 6 and 24 hours PR (hPR), as well as rRPE, in the presence of FGF2 at 6 hPR, when the cells are arrested in the cell cycle, and 24 hPR, when these cells are proliferating, according to our previous studies [11]. The cells for each condition were subjected to analysis by real time quantitative PCR (RT-qPCR) to evaluate mRNA levels relative to intact RPE. Our analysis showed that DNMTs, including *dnmt1*, *dnmt3a*, and *dnmt3b*, were significantly downregulated in the RPE at 6 hPR in the presence or absence of FGF2, while only *dnmt1* and *dnmt3b* were significantly downregulated in the RPE at 24 hPR in both conditions (Figure 2a). *Uhrf1*, an important factor for maintaining DNA methylation [50], was downregulated at 6 and 24 hPR in the t-rRPE (no FGF2) while it was upregulated in the rRPE (FGF2 present) at 24h (Figure 2a). Among the TET enzymes, only *tet3* was significantly upregulated at 6 and 24h PR in the rRPE as well as at 24 hPR in t-rRPE (Figure 2b). Similarly, DNA repair-associated factors *gadd45α,β,γ* [51], *prdm14*, a factor associated with hypomethylation in embryonic stem cells (ESC) and primordial germ cells [52] and thymine DNA glycosylase (*tdg*) were significantly upregulated at 6 and 24 hPR in t-rRPE and r-RPE (Figure 2b). We also analyzed the expression of components of the PRC2 *ezh2*, *suz12,* and *eed* (Figure 2c) and a group of Jumonji C (JmjC) domain-containing proteins involved in modulation of histone marks [53, 54], including: *jmjd1*, *jhdm1d*, *jmjd1c*, *jmjd4*, *jmjd5* and *kdm5* (Figure 2d). Among the components of PRC2, only *eed* was downregulated in the rRPE at 24 hPR (Figure 2c). In contrast, JmjC-containing proteins including *jmjd1* (also known as *kdm3a*), a histone lysine demethylase specific for H3K9me2/me1 that plays a role in transcriptional activation [55], was significantly upregulated at 24 hPR in the t-rRPE and both 6 and 24 hPR in the rRPE. *Jhdm1d* (also known as *kdm7a*, *kiaa1718*), a dual-specific histone demethylase specific for H3K9me2/me1 and H3K27me2/me1 [56], was downregulated only at 6 hPR in the t-rRPE. *Jmjd5* (also known as *kdm8*), a specific H3K36me2 lysine demethylase involved in cell cycle regulation and pluripotency in ES cells [57], was downregulated at 6h PR in the rRPE, and *kdm5b*, a H3K4me2/me3 demethylase [54], was downregulated at 24 hPR in the t-rRPE (Figure 2d). Our analysis did not show any significant changes in the expression of *utx* (*kdm6a*), a H3K27me3-specific demethylase [58]. Together, these results show that epigenetic factors associated with histone modifications, and moreover, DNA methylation and demethylation enzymes, including TET3, are differentially regulated in the t-rRPE and rRPE.

**Figure 2.**
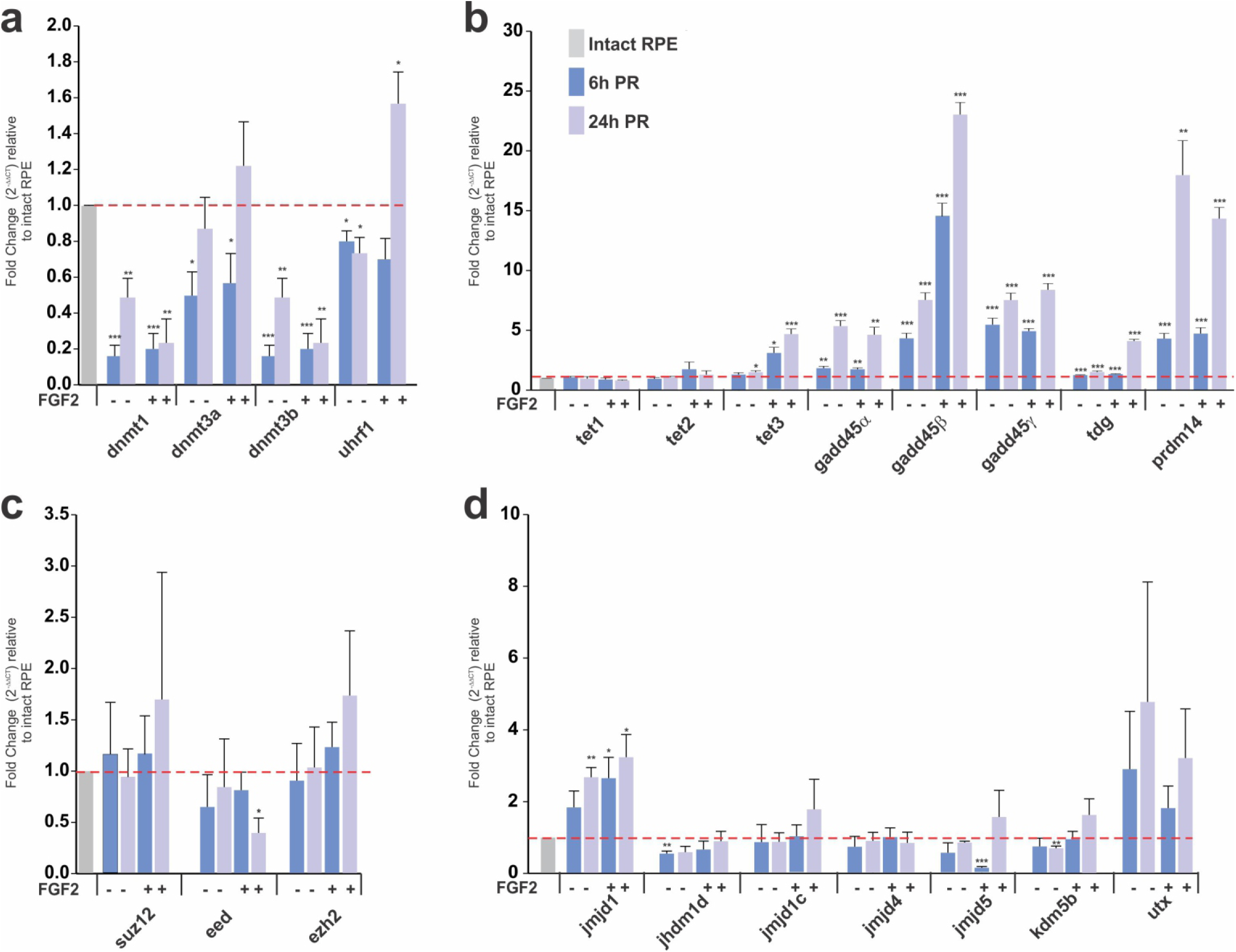
Differential expression of factors associated with histone and DNA modifications in the t-rRPE and rRPE. **(a-d)** Quantitative RT-PCR (qRT-PCR) analysis at 6 and 24h PR in the absence (t-rPE) or presence of FGF2 (rRPE). Relative expression of genes associated with **(a)** DNA methylation, **(b)** DNA demethylation and repair **(c)** components of PRC2 repressive complex and **(d)** histone lysine demethylases. Data are expressed as a Fold Change and were normalized with expression in intact RPE cells (dotted red line). Means ± standard error are shown. *=*P* < 0.05; **=*P* < 0.01; ***=*P* < 0.001. Unpaired Student’s *t* test, *n*= 3 biological samples were performed in triplicate.

### Early re-patterning and differential levels of histone and DNA modifications in the t-rRPE and rRPE

To investigate if the genome-wide distribution and levels of DNA modifications and bivalent chromatin (H3K27me3/H3K4me3) changes, during the Phase I of RPE reprogramming (Figure 3s), we performed immunofluorescence staining and high-resolution three-dimensional (HR-3D) reconstruction confocal microscopy of individual RPE nuclei. Our results showed that the levels of 5mC were significantly reduced in both t-rRPE and rRPE when compared to the intact RPE (Figure 3a-r and t). HR-3D analysis showed not only the signal intensity, but also subnuclear distribution patterns, indicating that 5mC was prominently dense in some regions of the nuclei and colocalized with heterochromatic areas detected as DAPI/5mC positive in the intact RPE (Figure 3d, j and p arrows). In contrast, 5mC density decreased during reprogramming suggesting some degree of hypomethylation, including within DAPI-intensive regions and chromocenters in the internal region of the nuclei (Figure 3e arrows and f). The oxidized form of cytosine, 5hmC, increased in the rRPE (Figure 3f and t); likewise, a significant increase was observed for 5caC in both t-rRPE and rRPE (Figure S1a-g).

**Figure 3.**
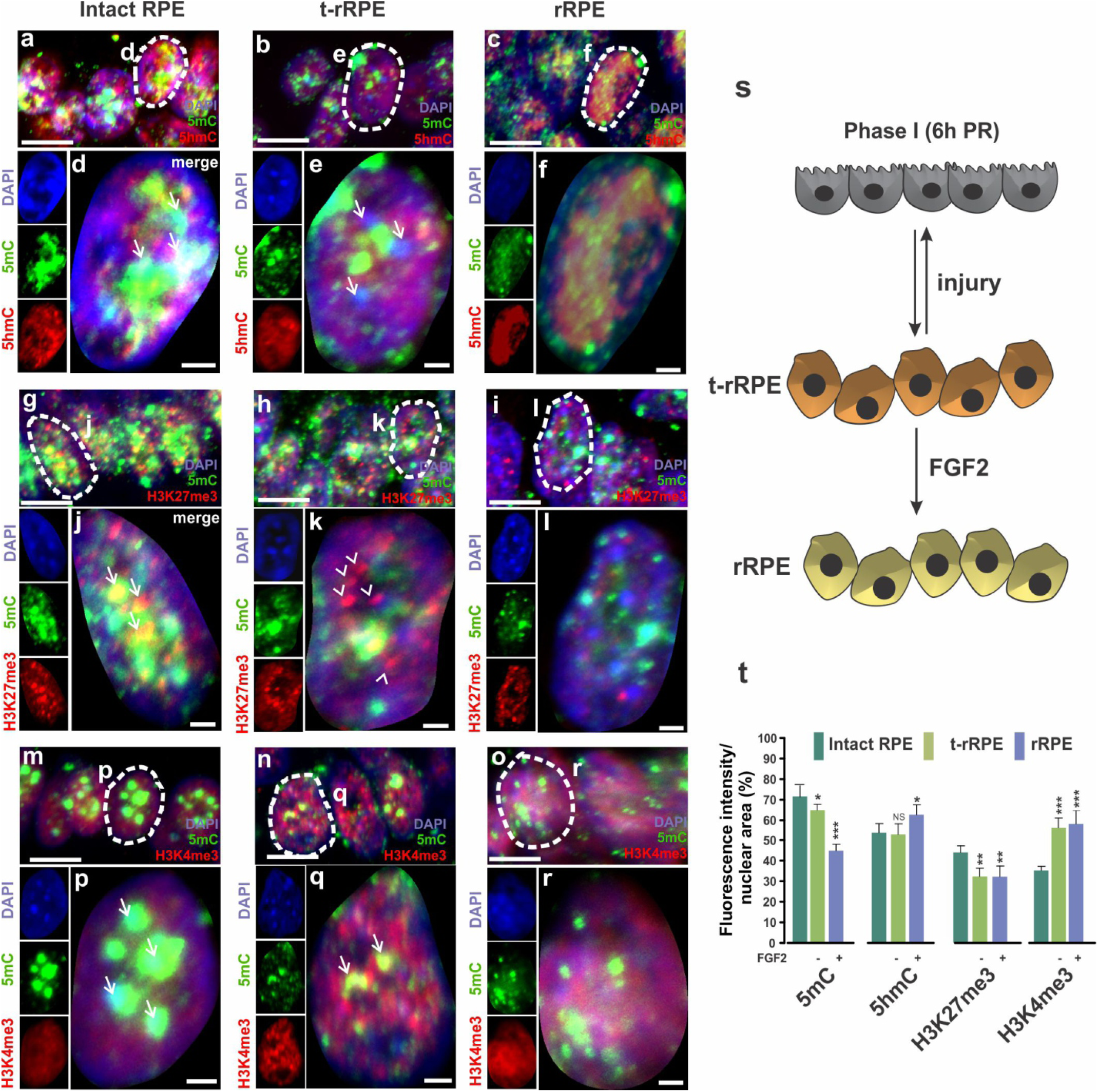
Early re-pattering and differential levels of histone and DNA modifications in t-rRPE and rRPE. Immunofluorescence staining and high-resolution three-dimensional (HR-3D) reconstruction confocal microscopy of histone modifications associated with bivalent chromatin (H3K27me3/H3K4me3) and DNA modifications (5mC and 5hmC) in **(a, g, m)** intact RPE at stage 23-25, **(b, h, n)** transient reprogrammed RPE (t-rRPE) and **(c, i, o)** reprogrammed RPE (rRPE) at 6 h PR. Single channels and merged views of the dotted areas are shown for each condition and combination of marks. Arrows in d, j and p indicate heterochromatic areas. Arrows in e show 5mC out of heterochomatic regions. Arrowheads in k show the marks don’t co-localize in t-rRPE. Arrows in q indicate co-localization of epigenetic marks. All images were processed in parallel and imaged using the same confocal settings. Scale bars in **a-c, g-i, m-o** are 5 μm. Scale bars in **d-f, j-l, p-r** are 1 μm. **(s)** Model of early RPE reprogramming (Phase I; 6 h PR). RPE is transiently reprogrammed (t-rRPE) in response to injury signals (e.g. retinectomy) and in the absence of FGF2. The reprogrammed RPE (rRPE) is generated in presence of FGF2. **(t)** Quantification of the percentage of fluorescence intensity per nuclear area of epigenetic marks in the t-rRPE or rRPE at 6 hPR, compared to intact RPE. *= *p*< 0.05, ***=*p*< 0.001. Student’s *t* test. NS, non-significant. Means ± standard error are shown.

The histone repression mark H3K27me3 was also present in heterochromatic DNA and co-localized with 5mC in dense DAPI regions of condensed chromatin in the intact RPE (Figure 3g, arrows in j). However, in the t-rRPE, H3K27me3 was dissociated from 5mC (Figure 3h and arrow heads in k) and was significantly reduced in the both t-rRPE and rRPE (Figure 3h-l and t). The activation mark H3K4me3 was distributed homogeneously throughout the nuclei and increased in the t-rRPE and rRPE (Figure 3m-r and t). We did not observe any significant changes in the activation mark H3K27Ac in t-rRPE and rRPE when compared to intact RPE (Figure S2a-g). All these results show that during the Phase I of RPE reprogramming, in early t-rRPE and rRPE, there is differential chromatin remodeling including a concomitant reduction of global DNA methylation toward a hypomethylated state and redistribution and changes in the levels of bivalent chromatin, suggesting gene activation (increased levels of H3K4me3 and reduced levels of H3K27me3). Moreover, these changes are independent of proliferation since at 6 hPR the t-rRPE and rRPE are still arrested in the cell cycle, as has been previously demonstrated [11]. The increased levels of 5hmC and 5caC in the rRPE and the induction of *tet3* mRNA suggest that DNA demethylation is involved in RPE reprogramming.

### Whole genome bisulfite sequencing reveals regions of differential DNA methylation in t-rRPE and early rRPE

To investigate whether the observed changes in DNA methylation occur at gene regulatory sequences, we performed WGBS in intact RPE, as well as in t-rRPE and rRPE at 6 hPR, using tissue isolated via LCM (see materials and methods). We then identified differentially methylated cytosines (DMCs), defined as CpG dinucleotides exhibiting methylation differences >20%, and DMRs, defined as genomic regions of differential methylation across multiple CpGs identified by BSmooth software (see methods, Supplementary Files 1-3, Figure S3). Between 8k–10k total DMRs were identified for each pairwise comparison, and analysis of the genomic distribution of the DMRs revealed a majority of changes in CpG methylation occurred proximal to coding regions (promoter /exon /intron) relative to intergenic regions (Figure 4a). Interestingly, when comparing intact RPE or t-rRPE to rRPE DMRs, more than half are hypomethylated (Figure 4a). We next turned our focus to changes in promoter region methylation, as past studies have revealed methylation of promoter regions is associated with transcriptional silencing [59]. Genes with a promoter region overlapping a DMR were extracted (Supplementary Files 4-6), and a total of 4,018 genes were identified together in intact RPE, t-rRPE and rRPE, with 239 genes containing a DMR in the promoter region in all 3 pairwise comparisons. Comparison of intact RPE to t-rRPE or rRPE revealed a disproportionally large number of shared genes (926), potentially reflecting common changes in DNA methylation as a response to injury alone or FGF2 exposure. Division of these genes into hypomethylated-only or hypermethylated-only DMRs revealed a similar shared enrichment (Figure 4b). In order to further visualize methylation patterns proximal to transcription start sites (TSSs), percent cytosine methylation surrounding the TSS (± 6000 bp) of all annotated genes was averaged, revealing global depletion of methylation at TSSs, accompanied by sharp increases in gene body methylation (Figure 4c), similar to what has been found in past studies [60]. Altogether, these results illustrate a dynamic DNA methylation landscape in response to retinectomy and FGF2, with a large proportion of DMRs occurring in or proximal to coding regions of the genome.

**Figure 4.**
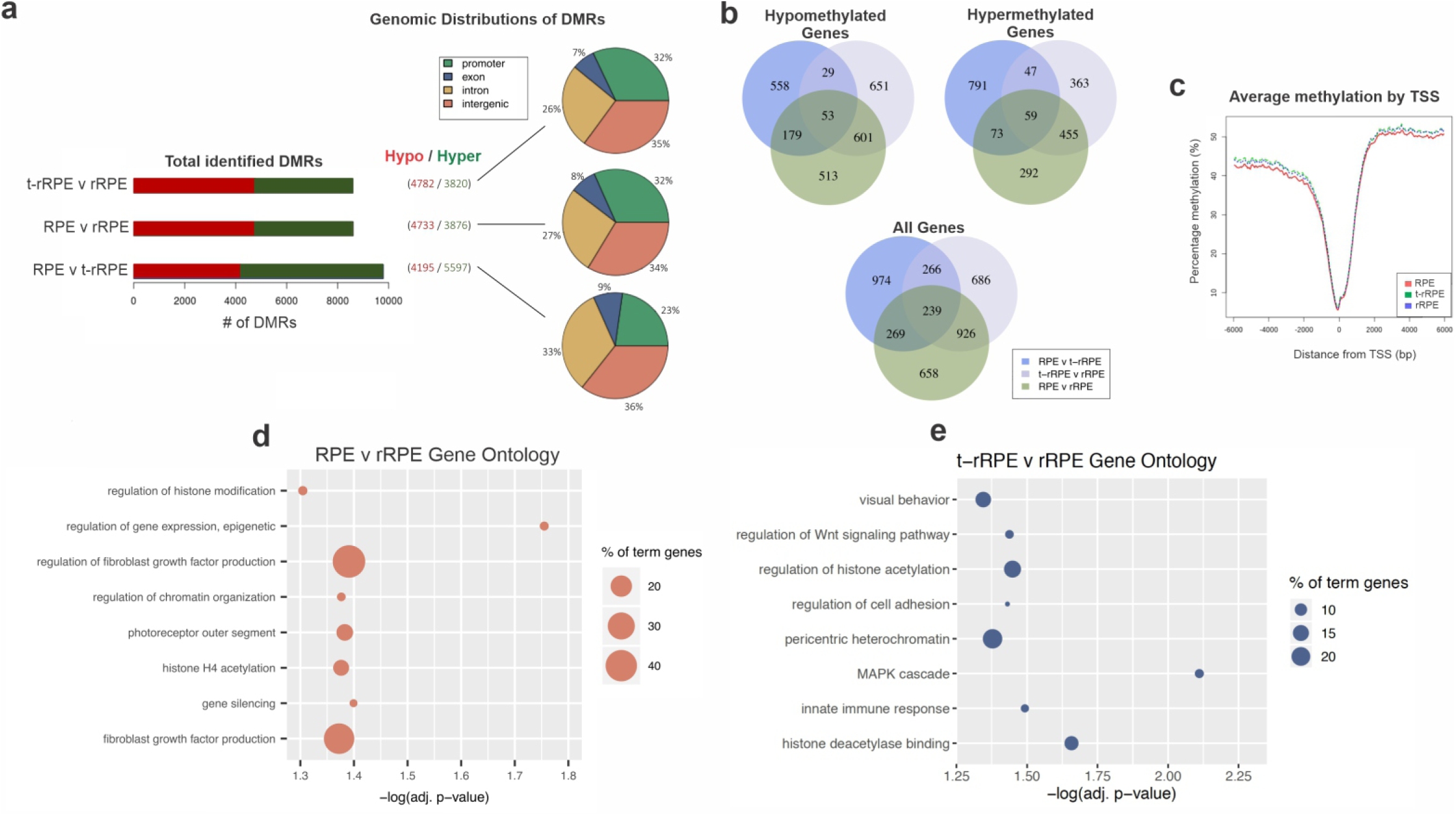
Whole genome bisulfite sequencing (WGBS) of intact RPE, t-rRPE, and rRPE reveal global changes in promoter region methylation. **(a)** Total identified DMRs, defined using the BSmooth software, are shown for each pairwise comparison in the context of hypo (red) or hypermethylation (green). Percentage of DMRs in context of their respective genomic features are shown in the pie chart. **(b)** Total genes with promoter region-containing DMRs for each pairwise comparison are shown in Venn diagrams, and broken down by hyper / hypo methylated genes. **(c)** Average percent methylation surrounding the transcription start site (TSS) is displayed for all annotated genes in the chicken genome. **(d, e)** Bubble charts display enriched Gene Ontology terms shown for RPE v t-rRPE and t-rRPE v rRPE pairwise comparisons, respectively. Y-axis indicates GO term, X-axis represents adjusted p-value, and bubble size corresponds to percentage of genes associated with each term identified in GO analysis.

### Loci-specific changes in DNA methylation occurs early during RPE reprogramming

To assign functionality to changes in DNA methylation at the promoter regions, we assessed our genes with identified DMRs for overrepresented gene ontology (GO) terms. Comparison of intact RPE and rRPE revealed significant enrichment in gene terms such as regulation of histone modification, epigenetic regulation of gene expression, and FGF production (Figure 4d), while comparison of t-rRPE and rRPE demonstrated enrichment in regulation of histone acetylation, MAPK cascade, and histone deacetylase binding, among others (Figure 4e). We have previously shown that activation of MAPK cascade is indispensable for RPE reprogramming, and its inhibition prohibits reprogramming even in the presence of FGF2 [61]. Comparison of RPE to t-rRPE did not yield significant GO results (Supplementary file 7). Full GO results, genes and networks can be found in the supplementary materials (Figure S4 and Supplementary Files 7-9). To further explore the role of methylation at the promoter region, we next turned our attention to methylation patterns found at individual promoter loci. Promoters of several genes upregulated in early RPE reprograming such as pluripotency inducing factors *sox2* and *lin-28a*, as well as EFTFs *six3*, *six6* (*optx2*) and *pax6* [11], were found in a hypomethylated state in the intact RPE as well as in t-rRPE and rRPE (Figure 5a and b). Interestingly, a distal region within the promoter of *lin-28a* displays hypomethylation in the rRPE compared to intact RPE and t-rRPE, however this sequence contains moderate methylation levels (< 40%) and it fails to reach statistical significance with our pipeline (Figure 5b). We have previously shown that *lin-28a* is upregulated in the presence of FGF2 (in rRPE), and *lin-28a* overexpression alone is capable of reprogramming RPE in the absence of FGF2 [11]. As such, the transcriptional responsiveness of *lin-28a* to FGF2 at embryonic day 4 may be reflective of its hypomethylated state. On the other hand, there were other groups of gene promoters that were identified to have hypomethylated DMRs in rRPE (Figure 5g), such as *trim35* (Figure 5c), indicating that these genes undergo changes in methylation in response to FGF2, highlighting *trim35* competence for transcriptional activation during reprogramming. Conversely, yet another group of gene promoters is represented by the *rho* promoter, which drives the expression of photoreceptor-specific gene rhodopsin. Genes like *rho* exist in a hypermethylated state throughout all conditions (Figure 5d). *Rho* is not expressed in the chicken retina until embryonic day 15, indicating methylation may play a role in its transcriptional repression during early RPE development and reprogramming [62]. The top 30 hypomethylated promoter regions for each pairwise comparison are displayed in heatmaps, indicating genes that undergo significant loss of methylation following injury alone (t-rRPE) or with FGF2 exposure (rRPE), and whose later activation may also play a role in the RPE-to-retina reprogramming process (Figure 5e-g). In conclusion, RPE displays methylation patterns that may underlie its reprogramming potential, given the low basal methylation of *sox2*, *lin-28a*, and key retinal genes, as well as site-directed changes in methylation in t-rRPE and rRPE.

**Figure 5.**
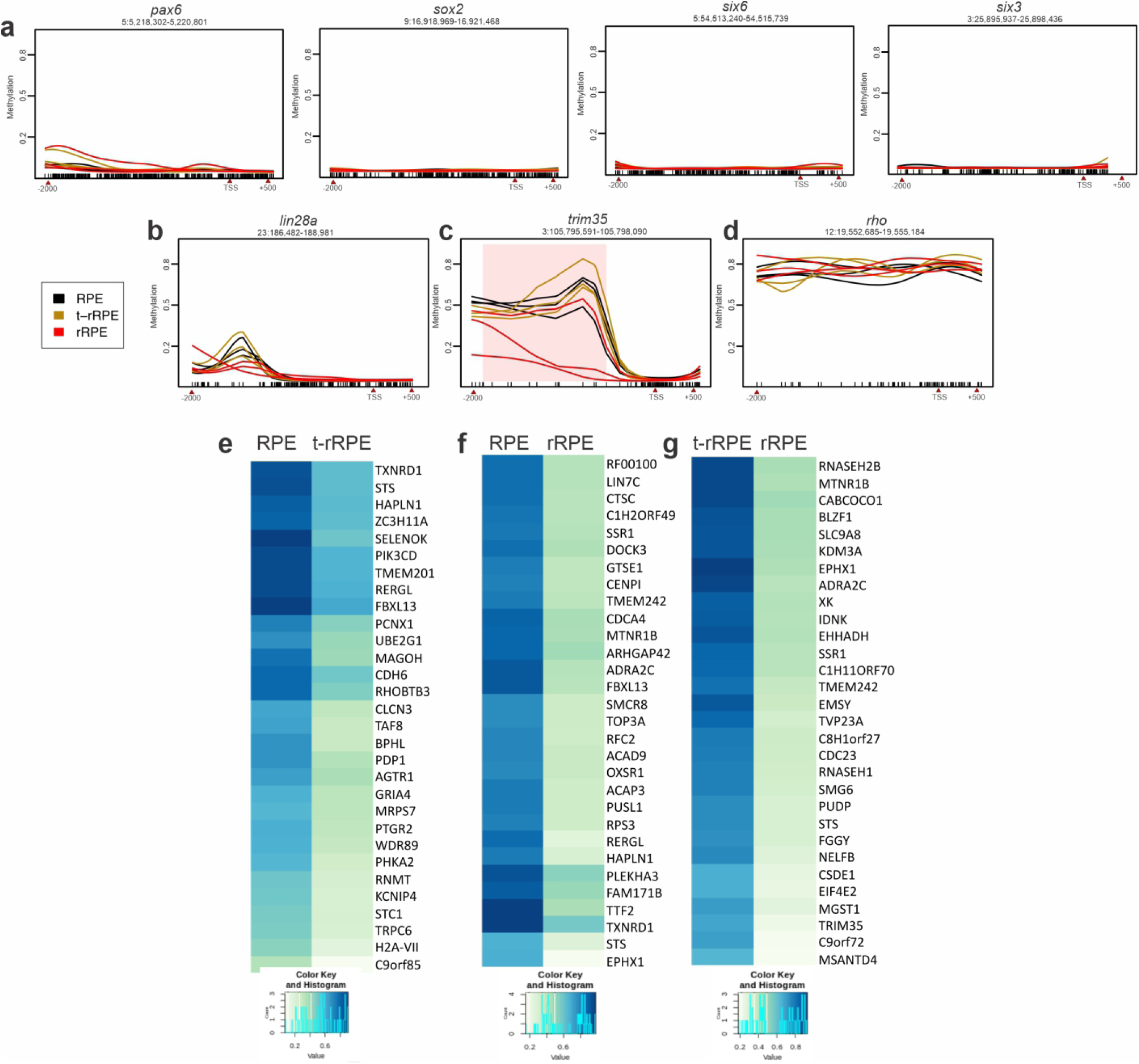
Whole genome bisulfite sequencing (WGBS) reveals methylation status of specific promoter regions. Pax6, sox2, six6, six3 **(a)**, lin28a **(b)**, trim35 **(c)**, and rho **(d)** promoter regions (−2000bp, +500bp from TSS) were plotted using smoothed methylation values generated by BSmooth. Y-axis indicates average methylation and X-axis indicates genomic coordinates, line color represents condition. DMR identified in trim35 is highlighted in pink. (**e,f,g**) Top 30 hypomethylated promoters, which display largest difference in average methylation, and corresponding methylation values for each pairwise comparison are illustrated in heatmaps.

### Active demethylation during reprogramming of RPE to NE

Given that we observed differential methylation and global reduction of 5mC simultaneously with increased levels of 5hmC in the early stage of rRPE, we further analyzed the levels of DNA modifications during the Phase II of RPE reprogramming (Figure 6a) in the newly formed NE that originated from the rRPE. Our results showed significant reduction of both 5mC and 5hmC in the NE in comparison to non-reprogrammed RPE (Figure 6b-j). Since TET enzymes can further oxidize 5hmC to 5fC and 5caC, we evaluated the levels of 5caC in the NE. The majority of non-reprogrammed RPE (pigmented cells) showed low levels of 5caC (Figure 6k-r). However, 5caC levels significantly increased and were more persistent among the cells present in the new NE (Figure 6k-r). The decreased levels of both 5mC and 5hmC, the increased levels of 5caC in the NE, as well as *tet3* upregulation at early times of reprogramming (Figure 2b), together suggest a state of hypomethylation mediated by active DNA demethylation that persists at 3 dPR into the new NE.

**Figure 6.**
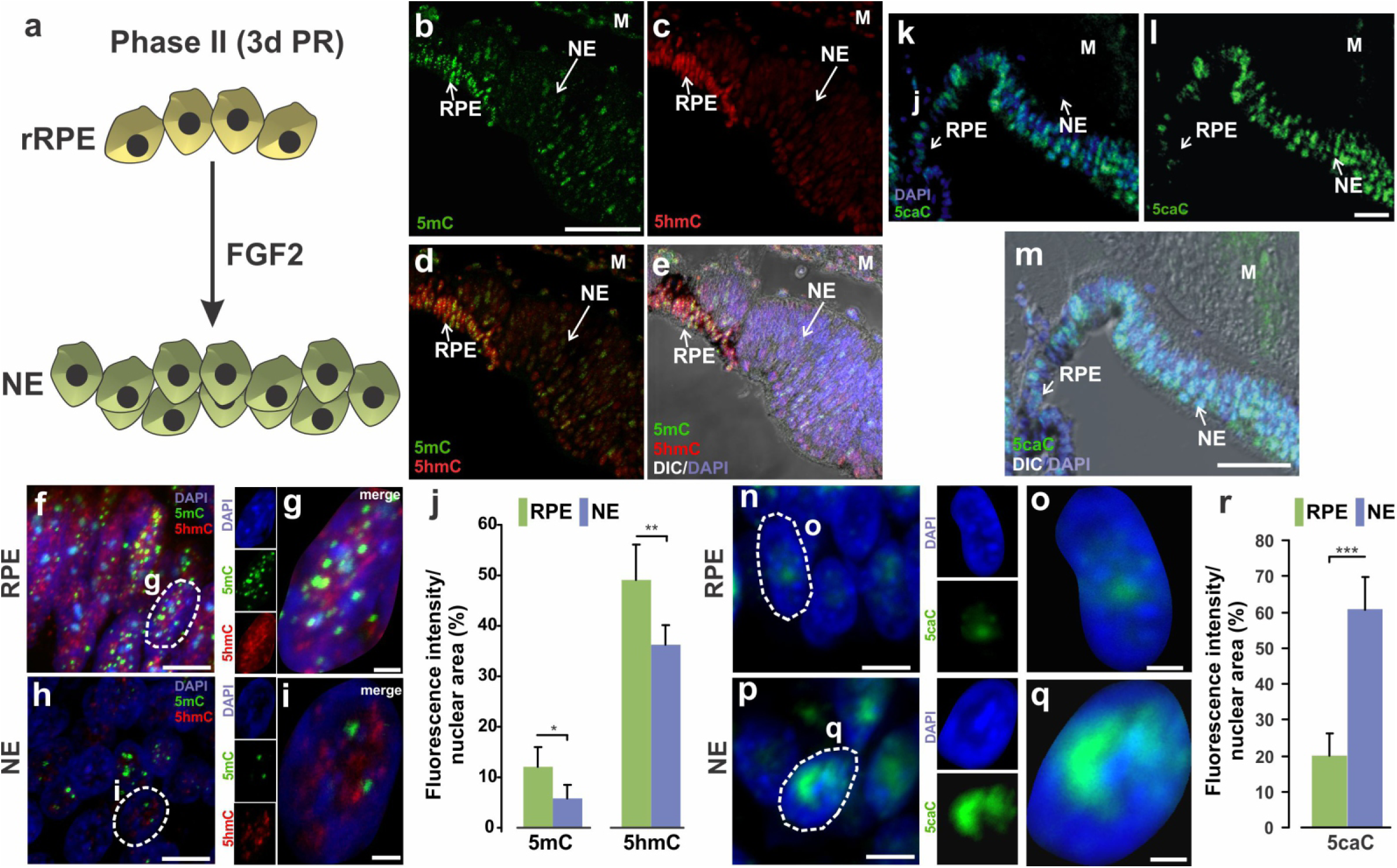
Dynamic changes of DNA modifications during RPE reprogramming to neuroepithelium (NE). **(a)** Model of RPE reprogramming to NE (Phase II). At 3d PR in presence of FGF2, proliferating rRPE give rise to retina progenitor cells that generate the new NE. **(b-i)** Immunofluorescence staining of 5mC (in green) and 5hmC (in red) at 3d PR. **(f-i)** High-resolution three-dimensional (HR-3D) reconstruction confocal microscopy images of RPE (pigmented area in **e**) and NE. **(j)** Percentage of fluorescence intensity per nuclear area for 5mC and 5hmC**. (k-q)** Immunofluorescence staining of 5caC at 3d PR. **(n-q)** HR-3D images of RPE (pigmented area in **m**) and NE. DAPI (blue), was used for nuclear counterstaining. Single channels and merged views of the marked areas with dotted lines are shown for each mark. All images were processed in parallel and imaged using the same confocal settings. Differential interference contrast (DIC) was used to illustrate the pigmented RPE in **e** and **m**. M: mesenchyme; RPE: Retinal Pigment Epithelium; NE: Neuroepithelium. Scale bar in: **b** is 50 um and applies to **c-e, l** is 50 um and applies to **k,** in **m** is 50 µm; in **f, h, n** and **p** is 5 µm; **g, i, o** and **q** is 1 µm. **(r)** Quantification of the percentage of fluorescence intensity per nuclear area for 5caC. *= *p*< 0.05, **=*p*< 0.01, ***=*p*< 0.001. Student’s *t* test. Means ± standard error are shown.

### Decreased levels of H3K27me3 correlates with increased levels of H3K27Ac in the new NE generated from rRPE

Due to the role histone modifications play during regeneration and reprogramming in other systems [41, 63, 64], we investigated if the patterns and levels of H3K27me3, H3K4me3, and H3K27Ac were modified at 3 dPR in the new NE originated from the rRPE. Concomitantly with the reduction of 5mC, H3K27me3 decreased significantly in the new NE compared with non-reprogrammed RPE (compare H3K27me3 signal in pigmented vs non-pigmented cells) (Figure 7b-f and s, Figure S5b-d). In contrast, our analysis did not show any significant changes in the levels of the activation mark H3K4me3 in the non-reprogrammed RPE cells and the new NE (Figure 7h-l and s, Figure S5f-h). We also observed a significant increase of the activation mark H3K27Ac in the NE compared to non-reprogrammed RPE (Figure 7n-s; Figure S5j-l). The dynamic changes of histone modifications observed at late stage of RPE reprogramming towards retina progenitors suggest significant changes in the bivalent chromatin by decreasing H3K27me3 and possible activation of distal regulatory sequences by enrichment of H3K27Ac [65].

**Figure 7.**
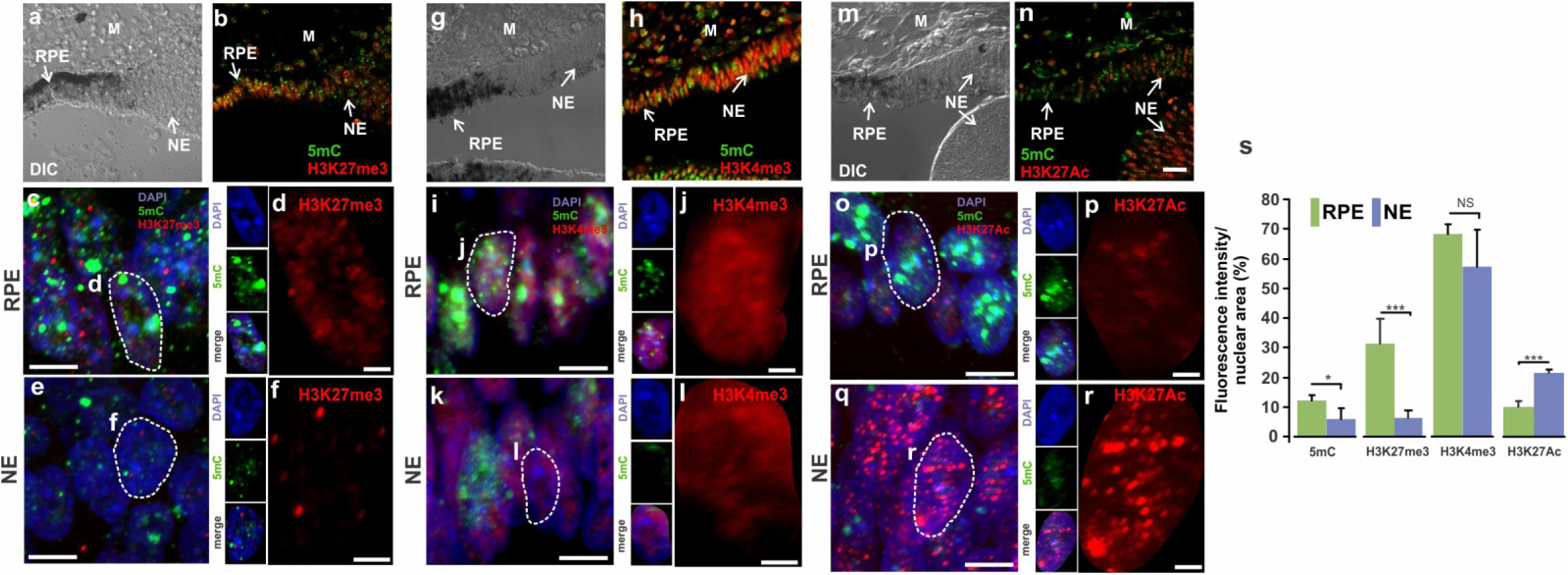
Re-patterning of histone modifications during RPE reprogramming to NE. Immunofluorescence staining and high-resolution three-dimensional (HR-3D) reconstruction confocal microscopy of histone modifications **(b-f)** H3K27me3 (red), **(h-l)** H3K4me3 (red), **(n-r)** H3K27Ac (red) along with 5mC (in green) at 3d PR in presence of FGF2. Single channels and merged views of the marked areas with dotted lines are shown for each combination of marks. HR-3D reconstruction confocal microscopy images of RPE, pigmented areas in **a**, **g** and **m**, are showed in **c**, **i** and **o**, respectively. All images were processed in parallel and imaged using the same confocal settings. Differential interference contrast (DIC) was used to illustrate the pigmented RPE in **a**, **g** and **m**. DAPI (blue), was used for nuclear counterstaining. M: mesenchyme; RPE: Retinal Pigment Epithelium; NE: neuroepithelium. Scale bar in **n** is 50 µm and applies to panels **a, b, g, h and m**. Scale bar in **c, e, i, k, o and q is** 5µm. Scale bar in **d, f, j, l, p** and **r** is 1µm. **(s)** Quantification of the percentage of fluorescence intensity per nuclear area of histone modifications and 5mC. *= *p*< 0.05, ***=*p*< 0.001. NS, non-significant. Student’s *t* test. Means ± standard error are shown.

### TET3 overexpression is sufficient to reprogram RPE to NE in the absence of exogenous FGF2

Our results show that during the Phase I of RPE reprogramming, at 6 hPR, 5mC significantly decreased while 5hmC increased in the rRPE (Figure 3t). In addition, *tet3* transcript was upregulated, and WGBS analysis showed differential methylation in the rRPE. Interestingly, the dynamic changes of DNA modifications at early times of rRPE occur independently of proliferation [11]. Later during Phase II, at 3 dPR when NE is generated from the proliferating rRPE [11], both 5mC and 5hmC decrease while 5caC increases. These results are compatible with a mechanism by which active elimination of global DNA methylation must occur before RPE reprogramming can proceed. To explore the role of TET3 in RPE reprogramming, we co-electroporated RPE 1 hPR with a plasmid containing the catalytic domain of mouse *TET3* (aa 697-1668) under the control of CMV promoter (pcDNA-Tet3, [66]) and a plasmid containing GFP (pIRES-GFP) as a reporter of electroporated areas (see materials and methods). Embryos collected 3 days post-electroporation showed prominent rRPE, determined by presence of NE tissue evaluated by histology (Figure 8b and f) and electroporated areas confirmed by GFP immunostaining (Figure 8e). In contrast, the histological analysis of RPE electroporated with control plasmids (pcDNA 3.1+pIRES-GFP) 1 hPR does not show NE tissue (Figure 8a, d and c). A systematic analysis of histological sections showed that in 50% (*n* = 18) of *TET3*-electroporated eyes, RPE was reprogrammed to NE (Figure 8c) and 38% show clear thickened and depigmented areas (not shown); this difference could be due to electroporation efficiency. In summary, these results strongly suggest that TET3 is sufficient to reprogram RPE to NE.

**Figure 8.**
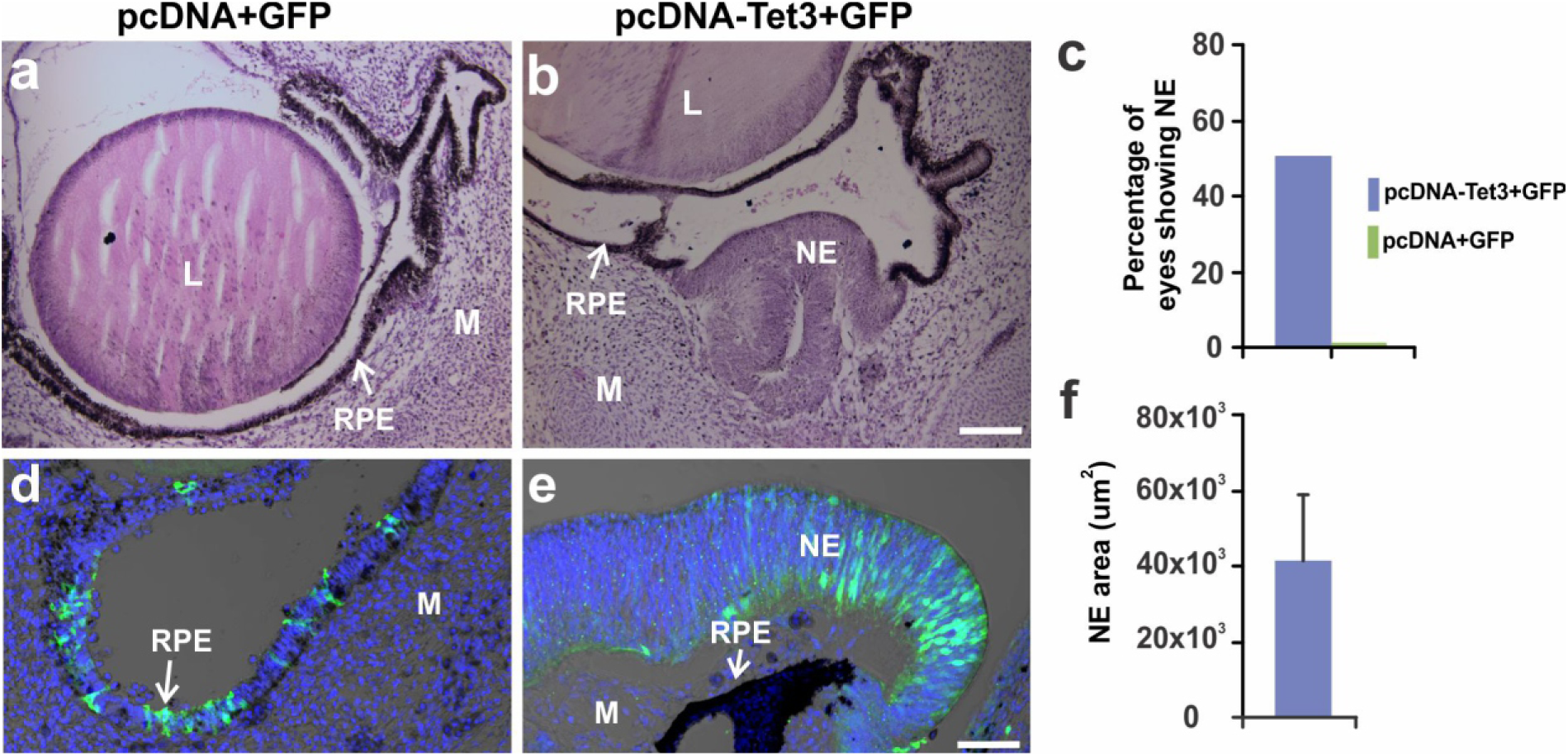
TET3 overexpression is sufficient to induce RPE reprogramming to NE. **(a,b)** Representative H&E histological sections at 3d PR and electroporation with **(a)** control plasmids pcDNA + GFP or **(b)** pcDNA-Tet3 + GFP (see material and methods). GFP immunofluorescence staining merged with DIC of electroporated eyes with **(d)** pcDNA + GFP or **(e)** pcDNA-Tet3 + GFP. L: lens; M: mesenchyme; RPE: retinal pigment epithelium; NE: neuroepithelium. Scale bar in **b** is 50 μm and applies to **a**. Scale bar in **e** is 20 μm and applies to d. **(c)** Percentage of eyes showing RPE reprogramming to NE at 3d PR after electroporation with pcDNA-Tet3 + GFP or pcDNA+GFP. DAPI (blue), was used for nuclear counterstaining. **(f)** Quantification of reprogrammed RPE in histological sections, expressed as NE area (µm^2^) at 3d PR after electroporation with pcDNA-Tet3+GFP. Mean ± standard error is shown.

## DISCUSSION

Previous studies have shown dynamic changes of histone modifications during mouse and human retina development [67–70], however, this is the first study to show dynamic changes in the levels of histone and DNA modifications during chick RPE reprogramming. It is established that chick RPE is competent to reprogram to neural retina only during a specific period of its early development (stages 23-25; embryonic day 4-4.5) [7, 8]. We therefore characterized the presence and distribution of histones and DNA modifications in the chick eye when the RPE is competent for reprogramming. Among the histone modifications analyzed, H3K27me3 was particularly enriched in the P-RGCs and reprogramming competent RPE (Figure 1h). Similar results were observed at Stage 27 (E5) and 31 (E7) (data not shown). During mouse retina development, H3K27me3 is enriched in the inner neuroblastic layer (INBL) and GCL at E16, E18, P0 (when RGCs begin to differentiate) and postnatal day 1 (PN1) [67, 69], suggesting that H3K27me3 may be associated with neural retina differentiation in post-mitotic neurons. Our results here suggest that the chick embryo recapitulates the patterns of H3K27me3 observed during mouse retina development particularly during RGCs differentiation. Also, the accumulation of H3K27me3 observed in the reprogramming competent RPE could be due to its post-mitotic state as it occurs in cells from other tissues including mouse and *xenopus* retina as well as in chick spinal cord [67, 71, 72]. However, we cannot rule out the possibility that the accumulation of H3K27me3 in the reprogramming competent RPE can be related to the bivalent state of the chromatin since we also observed high levels of H3K4me3. Interestingly, a recent report showed that the chromatin state of murine adult (2.5-3 month-old) RPE contains several key genes for RGCs, bipolar, amacrine, and horizontal cells marked with PCR2 repressive mark H3K27me3 [45].

In addition to P-RGCs and the reprogramming competent RPE, H3K27me3 is present in the most anterior part of the eye that contains multipotent optic cup stem/progenitor cells (Figure 1a). In *Xenopus* eye at Stage 41 (tadpole stage), transcripts of the components of the PRC2 complex including *ezh2*, *suz12*, *eed*, and *rbbp4* are present in the undifferentiated proliferating cells present in the ciliary margin zone (CMZ) [72]. It is possible that the genes regulated by H3K27me3 in the different species may differ between the post-mitotic differentiated retina cells, the undifferentiated multipotent progenitors present in CMZ and the reprogramming competent RPE. Therefore, further Chromatin Immunoprecipitation Sequencing (ChIP-seq) experiments will help to elucidate the genes associated with H3K27me3 and H3K4me3 in the reprogramming competent RPE. Our results also demonstrated that the reprogramming competent RPE, the P-RGCs, and the CM are enriched in 5hmC (Figure 1f, m). Similar results were observed in the P-RGCs and differentiated RGC at Stages 27 and 31, respectively (data not shown). In mouse embryonic stem (ES) cells, 5hmC is highly enriched at transcription start sites (TSSs) of promoters that contain bivalent chromatin (H3K27me3/H3K4me3) [73]. In adult mice, 5hmC is particularly abundant in the central nervous system (CNS) [74, 75] and is associated with neural development and differentiation [76–78]. However, in the context of eye and retina development, limited information is available concerning the presence of 5hmC. In mice at week 2 (eye opening) and week 3 (mature state), 5hmC is accumulated in euchromatic regions suggesting its association with transcriptional activity in RGCs and the inner nuclear layer (INL) [79]. In addition, genes enriched in 5hmC have a positive correlation with active transcription during retina maturation [79]. Another study showed the accumulation of 5hmC in post-mitotic mouse neurons associated with active transcription and in functional demethylation by decreasing the occupancy of methylcytosine binding protein MeCP2 in the transcript unit, and in consequence activating gene expression [80]. To our knowledge, this is the first time that 5hmC is detected in the embryonic RPE and CM that host the multipotent optic cup stem/progenitor cells, the two sources of cells for retina regeneration in the embryonic chick [10, 81]. Analysis of the genes associated with 5hmC by hydroxymethylated DNA immunoprecipitation (hMeDIP) or TET-Assisted Bisulfite Sequencing (TAB-Seq) could help to elucidate the genomic distribution, and the target genes associated with 5hmC and H3K27me3 in the reprogramming competent RPE. The fact that H3K27me3 and 5hmC show similar patterns at later stages of retina development (27 and 31) (when the RPE losses its plasticity to regenerate the retina) suggests that these epigenetic marks in the NE are more associated with a developmental and differentiation program of the retina. Supporting this idea, it has been shown that global levels of 5hmC and H3K27me3 are highly correlated in multiple tissues, and their pattern and distribution are associated with differentiation [82].

We previously reported that at early times after retina removal, the reprogramming competent and quiescent (p27^Kip1^+/BrdU-) RPE is transiently reprogrammed (t-rRPE) in response to injury (retinectomy) (Figure 3s). In the presence of an exogenous source of FGF2, the t-rRPE can be further reprogrammed (rRPE, Figure 3s) to produce retina progenitors that eventually proliferate generating a new NE (Figure 6a) [11]. Here we show that the levels of 5mC and bivalent chromatin (H3K27me3/H3K4me3) are differentially regulated at 6 hPR in the t-rRPE in absence of proliferation. These results indicate that removal of repressive marks 5mC and H3K27me3 as well as injury-induced changes of H3K4me3, which is associated with active promoters and hypomethylated promoters are the first step for the initial reprogramming of RPE. In accordance with the reduction of 5mC, we observed significant downregulation of DNMTs and the critical regulator of DNA methylation *uhrf1*, a factor that is associated with liver regeneration and redistribution of H3K27me3 from promoters to transposons [83]. Our results obtained at 6 hPR in t-rRPE, partially recapitulate the results observed during fin regeneration in zebrafish where the levels of 5mC and 5hmC were transiently reduced at 24 hours post-amputation (hpa) in the blastema cells, and such reduction was also independent of proliferation [37]. However, in the t-rRPE we did not detect significant changes in the levels of 5hmC evaluated by immunofluorescence staining and only small but significant upregulation of *tet3* was observed at 24 hPR (Figure 2b). In addition, our results demonstrated an increase in the levels of 5caC (Figure S1) and upregulation of genes associated with DNA repair and *tdg* (Figure 2b) suggesting a process of active demethylation in the t-rRPE. One possible explanation for the discrepancy between the levels of 5hmC and 5caC is the rapid oxidation of 5hmC or the accumulation of 5caC as has been demonstrated during linage specification of neural stem cells (NSCs) [84]. In line with our results, during Müller Glia (MG) reprogramming in zebrafish, changes in DNA methylation and a correlation between promoter DNA demethylation were observed at early times after injury [35]. These results suggest that dynamic changes in DNA methylation may be a common mechanism in MG and RPE reprogramming. In contrast to our results, during MG reprogramming, *dnmt3* and *dnmt4* were moderately increased at 15 hours post-injury (hpi) while *dnmt1*, *dnmt4*, *dnmt5,* and *dnmt7* were upregulated significantly in Müller Glia progenitor cells (MGPC) at 4 days post-injury (dpi) [35, 40]. These differences suggest that, in contrast to the zebrafish MG reprogramming, during transient chick RPE reprogramming there is an impaired process of DNA methylation, followed by a process of demethylation; however, we cannot rule out the possibility that such processes occur concomitantly at early times PR. In agreement with this, it has been shown that DNA demethylation is associated with the process of cellular reprogramming [85]. Meanwhile, our WGBS results demonstrated that the promoters associated with pluripotency inducing factors such as *sox2* and *lin28a* as well as EFTFs and retina progenitor markers such as *six3*, *six6* and *pax6* were already hypomethylated in the intact RPE. These results highlight some similarities between MG and RPE reprogramming in terms of DNA methylation, where basal hypomethylation is present in genes associated with regeneration and reprogramming. Interestingly, two independent reports from the same group suggested that the methylome of adult mice RPE and MG have similar methylation patterns to retinal progenitor cells (RPCs); these observations support the possibility that DNA methylation patterns are retained and demethylation may be one of the most important drivers for RPE reprogramming [45, 46]. Moreover, in adult mice after NMDA-induced retinal injury, *oct4*, the key pluripotency inducing factor, was transiently expressed in MG, but its expression was not sustained after 24h post-injury. In the same study, intravitreal administration of SGI-1027, a DNA-methyltransferase inhibitor, was able to induce and sustain the expression of *oct4*, suggesting that DNA methylation may be the first barrier that prevents adult mice MG from acquiring multipotency [36]. In line with the idea that demethylation may be one of the important drivers for reprogramming, we observed a significant increase of 5hmC and 5caC as well as the induction of *tet3* mRNA in the rRPE at 6 hPR in absence of proliferation, suggesting active demethylation independently of replication. Importantly, in agreement with zebrafish fin regeneration [37], we observed decreased levels of 5mC and 5hmC while a significant increase of 5caC at 3 dPR in the new NE generated from the rRPE. These results are consistent with a model of demethylation during RPE reprogramming to retina progenitors. However, it is also possible that the reduction of 5hmC is replication-dependent dilution since FGF2 induces proliferation at 3 dPR in the NE [11, 81]. Our results strongly suggest that a demethylation program initiated by injury is stimulated and sustained by FGF2, and this may link exogenous signaling with changes in DNA methylation, chromatin configuration, and ultimately gene expression. However, the relationship between FGF signaling and TET-mediated demethylation remains to be explored.

The relationship between 5hmC and histone modifications is complicated. 5hmC is highly enriched at gene promoters associated with bivalent chromatin, promoters with intermediate CpG density and CpG islands with low to medium GC-content [73, 86–88]. In mouse ESC, TET1 forms a functional complex with PRC2 at H3K27me3 marked regions and TET proteins contribute to epigenetic plasticity during cell differentiation [89, 90]. We observed that in the t-rRPE there was a decrease in 5mC and H3K27me3 and an increase of H3K4me3. Unexpectedly, we did not observe significant changes in the levels of expression of most of the components of PRC2 except for *eed* which was downregulated at 24 hPR in the presence of FGF2. Interestingly, during Schwann cell reprogramming in peripheral nerve injury, knock-out of the EED subunit of PRC2 leads to de-repression of genes associated with injury response by decreasing levels of H3K27me3 and increasing levels of H3K4me3 at promoter regions [91, 92].

The fact that *tet3* is induced during MG reprogramming at 4 days post-injury [35] and in the t-rRPE and rRPE, clearly suggest that TET3 may be a prime factor involved in the process of demethylation. In this regard, TET1, TET2, and TET3 catalyze the oxidation of 5mC to 5hmC, however they seem to have specific functions in different cellular contexts. TET1-2 are important for ESC linage specification [93, 94]. TET1 plays an important role during DNA demethylation and gene expression in mouse ES cells [86–88, 95] while TET3 contributes to genome-wide DNA demethylation and can efficiently reprogram fibroblast directly into functional neurons and induce demethylation of Ascl1, an important factor for MG reprogramming in mouse and zebrafish [96–102]. Moreover, TET3 is required for significant axon regeneration in the dorsal root ganglion (DRG) neurons, suggesting that an epigenetic barrier can be removed by active DNA demethylation mediated by TET3 [103]. In *Xenopus*, TET3 plays an essential role in early eye and neural development, and its depletion causes reduction of EFTFs *pax6*, *rx*, and *six3*; neural markers *ngn2* and *tubb2b*; and neural crest markers *sox9* and *snail* [104]. The zebrafish genome encodes single well-conserved orthologs of *tet1*, *tet2,* and *tet3,* and a recent study demonstrated that TET2 and TET3 are required for retinal neurogenesis [105]. For the first time, we demonstrated that TET3 is sufficient to induce RPE reprogramming to NE by 3 dPR in absence of exogenous FGF2. Interestingly, the overexpression of TET3 in the reprogramming competent RPE (uninjured eye) was not enough to induce reprogramming (data not shown), suggesting that injury signals, along with concomitant resolution of bivalent chromatin, are required in the first step of RPE reprogramming (t-rRPE) and the progression of a demethylation program allows for sustain reprograming (rRPE) that can form a NE and eventually a differentiated retina. In conclusion, we demonstrate that during the process of RPE reprogramming, injury signals induce dynamic changes in the chromatin including bivalent domains and DNA methylation/demethylation. The relationship between RPE reprogramming and DNA methylation has been established, and we for the first time identify TET3 as an important factor for RPE reprogramming. The analysis of the epigenetic landscape during regeneration is in its infancy; further experiments including ChIP-seq of histones modifications or high-resolution chromatin profiling from limited number of cells using CUT&RUN [106], as well as TET3-ChIP-seq, Assay for Transposase-Accessible Chromatin (ATAC)-seq, and Assisted Bisulfite Sequencing (TAB-Seq) to identify 5hmC on specific loci will provide comprehensive information of the epigenetic landscape of RPE reprogramming towards retina regeneration.

---

**Figure S1. 5caC is significantly increased at early times of t-rRPE and rRPE.** Immunofluorescence staining and high-resolution three-dimensional (HR-3D) reconstruction confocal microscopy of 5caC in **(a,d)** intact RPE at St 23-25 **(b,e)** transient reprogrammed RPE (t-rRPE) and **(c,f)** reprogrammed RPE (rRPE) at 6h PR. Single channels and merged views of the marked areas with dotted lines are shown and were processed in parallel and imaged using the same settings. DAPI (blue), was used for nuclear counterstaining. Scale bars in **a-c** are 5 µm. Scale bars in **d-f** are 1 µm. **(g)** Quantification of the percentage of fluorescence intensity per nuclear area for 5caC. L: lens. **=p< 0.01, ***=*p*< 0.001. Student’s *t* test. Means ± standard error are shown.

**Figure S2. Analysis of 5mC and activation mark H3K27Ac during early times of t-rRPE and rRPE.** Immunofluorescence staining and high-resolution three-dimensional (HR-3D) reconstruction confocal microscopy of 5mC (green) and H3K27Ac (red) in **(a,d)** intact RPE at St 23-25 **(b,e)** transient reprogrammed RPE (t-rRPE) and **(c,f)** reprogrammed RPE (rRPE) at 6h PR. Single channels and merged views of the marked areas with dotted lines are shown and were processed in parallel and imaged using the same confocal settings. Scale bars in **a-c** are 5 µm. Scale bars in **d-f** are 1 µm. **(g)** Quantification of the percentage of fluorescence intensity per nuclear area for 5mC and H3K27Ac. *= *p*< 0.05, ***=*p*< 0.001. NS, non-significant, Student’s *t* test. Means ± standard error are shown.

**Figure S3. Whole genome bisulfite sequencing data processing pipeline.** Raw reads were trimmed of adapters and low-quality bases using Trim Galore before mapping to Gallus-gallus-5.0 genome assembly and extracting methylation using Bismark. Resultant methylation count tables were used for DMC and DMR identification using MethylKit and BSmooth, respectively.

**Figure S4. Gene ontology networks produced by ClueGO.** Promoter region DMRs were identified and searched for overrepresented Gene Ontology terms using the Cytoscape plug-ins ClueGO and AutoAnnotate. **(a)** Network displaying enriched terms found in comparison of RPE and rRPE, and **(b)** network of terms produced by comparison of t-rRPE and rRPE. Enriched ClueGO term groupings are indicated by node color, and significance is represented by node size. Edges between nodes indicate term relationship. Only GO terms with p < 0.05 are depicted.

**Figure S5. Repatterning of histone modifications and 5mC during reprogramming of RPE to NE.** Immunofluorescence staining of histone modifications **(c-d)** H3K27me3 (red), **(g-h)** H3K4me3 (red), **(k-l)** H3K27Ac (red) and **(b, f, j)** 5mC (in green) at 3d PR in presence of FGF2. Merged views are shown in **d**, **h** and **l**. Differential interference contrast (DIC) was used to illustrate the pigmented RPE in **a**, **e** and **i**. DAPI (blue), was used for nuclear counterstaining in **d**, **h** and **l**. Scale bar in **l** is 50 µm and applies to all panels. L: lens; M: mesenchyme; RPE: retinal pigment epithelium; NE: Neuroepithelium.

## METHODS

### Chick embryos and surgical procedures

White Leghorn chicken eggs from Michigan State University were incubated in a humidified rotating incubator at 38°C. Retinectomies and FGF2 treatments were performed at embryonic day 4 (Stages 23-25, [6]) as previously described [10, 11, 61]. Embryos were collected at 6 and 24h or 3 dPR and processed for histology, immunofluorescence staining, or laser capture microdissection.

### Laser capture microdissection (LCM)

LCM, RNA extraction, and SPIA cDNA preparation were performed as previously described [11]. Briefly, intact embryos at St 23-25 or 6 and 24h PR were collected, infiltrated with different concentrations of sucrose and embedded in 2:1 25% sucrose: OCT compound (Sakura Finetek, Torrance, CA). Cryosections (12 µm) were collected on PEN Membrane Frame Slides (Thermo Fisher Scientific, LCM0521), followed by hematoxylin staining (Sigma, St. Louis, MO). LCM was performed using Veritas Laser Capture Microdissection system (Molecular Devices) to collect RNA using the settings previously described [11] or Arcturus XT (Life Technologies Carlsbad, CA) to collect DNA samples. DNA extraction was performed using PicoPure DNA Extraction Kit (Thermo Fisher, Waltham, MA), quantified using Quant-iT dsDNA Assay Kit high sensitivity (Thermo Fisher Scientific, Waltham, MA) or NanoDrop3300 and used in WGBS experiments.

### RT-qPCR

RT-qPCR was performed in a 20µl reaction containing 5μl (4ng) of SPIA cDNA, 10μl of 2x SYBR Green/Fluorescein qPCR Master Mix (SABiosciences, Qiagen, Maryland, USA) and primer mix to a final concentration of 500nM. All reactions were performed in triplicate using iCycler (BioRad, Hercules, CA) and amplification conditions 10 min at 95°C, 35 cycles of 15 sec at 95°C and 1 min at 60°C, followed by 65°C to 95°C for melting curve analysis. Splice junction primers were designed using Primer 3 (v 4.0) [107] and were optimized following the guidelines described in [108]. Primers sequences and Ensembl or GenBank ID are described in Table S2. Comparative cycle at threshold (2^-ΔΔCt^) and unpaired Student’s *t*-test analysis were used to determine relative changes in transcript levels compared to *gapdh* mRNA levels as previously reported [109]. All analyses were performed in triplicate with at least three independent biological replicates.

### Primary and Secondary Antibody reagents

Primary antibodies details and dilutions include, rabbit anti-H3K27me3 (Millipore, 07-449, 1:200), rabbit anti-H3K4me3 (Millipore, 07-473, 1:200), rabbit anti-H3K4me2 (Millipore, 07-030, 1:500), rabbit anti-H3K27Ac (Active Motif, 39133, 1:500), mouse anti-5mC (33D3) (Abcam, ab10805, 1:100), rabbit anti-5hmC (Active Motif, 39791, 1:200), rabbit anti-5caC (Active Motif, 61225, 1:3000, including TSA amplification), rabbit anti-GFP (Abcam, ab6673, 1:100), rabbit anti-HRP antibody (PerkinElmer, NEF812001EA, 1:3000), goat anti-rabbit Alexa Fluor-546 (ThermoFisher Scientific, A-11035, 1:100), goat anti-mouse Alexa Fluor-488 (ThermoFisher Scientific, A32723, 1:100), donkey anti-rabbit Alexa Fluor-488 (ThermoFisher Scientific, A32790, 1:100).

### Immunohistochemistry

Embryos were fixed in 10% Neutral Buffered Formalin (Thermo Scientific, Kalamazoo, MI) overnight (ON) at 4°C, transferred to 70% ethanol, embedded in paraffin, sectioned at 12 µm, and deparaffinized. Briefly, following antigen retrieval (0.01M Sodium Citrate, pH=6.0, 15 min), permeabilization (1% saponin, 5 min) (Sigma, St. Louis, MO) and blocking with 10% normal goat or donkey serum (Sigma, St. Louis, MO) in PBST (0.1% Triton-X100 in PBS) during 30 min at 37 °C, primary antibodies were diluted in blocking solution as indicated in the antibodies section and incubated ON at 4 °C. After washing in PBST (3 × 5 min), the samples were incubated 1h in the dark at room temperature (RT) with the corresponding secondary antibody (typically 1:100 in blocking solution) and DAPI 30 nM in PBS (Invitrogen, USA Grand Island, NY) for nuclear counterstaining was included before cover-slipped using Fluoromount (Sigma, St. Louis, MO). Controls to demonstrate the specificity of the antibody were included for every experiment, including samples with secondary antibody in the absence of primary. 5caC immunofluorescence staining was performed using a Tyramide Signal Amplification system (TSA plus Fluorescein, NEL741001KT, PerkinElmer), following the manufacturer’s instructions, including quenching of endogenous peroxidase for 20 min with 1% H_2_O_2_ and introduction of rabbit anti-HRP antibody ON at 4°C. All TSA amplifications were performed during 10 min at RT and negative controls no amplified or amplified (no primary antibody added), were included and used during confocal imaging.

### Confocal microscopy and image analysis

Confocal images (size 1024 × 1024) were collected on a Zeiss 710 Laser Scanning Confocal System (Carl Zeiss, Göttingen, Germany) using a x20/0.80 NA WD=0.55 or EC Plan-Neofluar 63×/0.75 Corr M27 objective lenses and a digital zoom of 7 to capture nuclei images. Imaging of each 488, 546 and DAPI were acquired sequentially in order to avoid bleed-through and pinhole size was consistently 1.0 airy unit (AU). Hight resolution images were obtained by collecting serial optical sections at increments of 0.3 µm apart in Z-stacks of about 15 images and processed with ZEN 2012 Browser (Carl Zeiss, Göttingen, Germany). Three-dimensional reconstruction (3D) of 2D confocal imaging was performed as previously described in [84]. Signal intensity analysis and quantification were performed using Image J Fiji plugin [110]. Briefly, each individual nucleus was selected using the freeform drawing tool in ZEN 2012 Browser and then processed. The positive immunofluorescence staining areas for each epigenetic mark and the associated mean fluorescence were measured by creating a binary image. The values are expressed as a percentage of epigenetic mark relative to DAPI from five different biological replicates in each experimental group. An unpaired Student’s *t*-test was applied for mean comparison using SigmaPlot 8.0 Software.

### *In ovo* electroporation

*In ovo* electroporations were performed as previously described [11, 111]. Briefly, using a Pico-injector system PLI-100 (Harvard Apparatus, Holliston, MA) and glass capillary needles, embryos were injected 1 hPR with 3μl of a mixture of pcDNA-Flag-Tet3 (3µg) (Addgene Massachusetts, USA, plasmid 60940, [66] and pIRES-GFP (1µg), or 3μl of a mixture at the same concentration of Flag-HA-pcDNA-3.1 (Addgene Massachusetts, USA, plasmid 52535) and pIRES-GFP as controls. Electroporated embryos were returned to the incubator and collected 72h post-electroporation and processed for immunohistochemistry and histology.

### Histology and quantification

Embryos were fixed in Bouin’s fixative (Ricca Chem. Comp., Arlington, TX) overnight at room temperature, embedded in paraffin and sectioned at 12µm. H&E stained sections were photographed with an Olympus BX51-P microscope with Magnifire S99800 Camera (Olympus, Tokio, Japan) and the images were processed and analyzed using Image J [112] to quantify the NE generated from rRPE.

### Whole Genome Bisulfite Sequencing (WGBS) and Data Processing

Genomic DNA (gDNA) samples isolated via LCM were sent to the University of Michigan Epigenomics core for library preparation and sequencing. The gDNA samples were quantified using the Qubit broad range dsDNA kit, and quality was assessed with Agilent’s genomic DNA 2200 TapeStation kit. A total of 100ng was used for each WGBS library preparation. Each sample was spiked-in with 0.5% (w/w) of unmethylated Lambda DNA prior to library preparation, according to the ENCODE consortium’s guidelines. DNA was fragmented to a range of 200-1000bp on a Covaris S220 focused-ultrasonicator system. The samples were next processed for end-repair and A-tailing, before ligation of indexed methylated duplex adapters. The ligation products were cleaned with two rounds of AMPure XP beads before running on a 1% agarose gel for selection of fragments between 200-650bp. After gel extraction, the ligated products were submitted to bisulfite treatment using Zymo’s EZ DNA Methylation kit, with the following protocol: 55 cycles of 95C for 30sec, 55C for 15 minutes. Samples were kept at 4C until cleanup. Final library amplification was done using the Roche FastStart Hi-fidelity DNA polymerase system. 16 cycles of amplification were used. The PCR products were then cleaned with AMPure XP beads, and the final libraries were quantitated using the Qubit High Sensitivity dsDNA kit. Quality of the libraries was assessed on Agilent’s 2200 TapeStation, using the High Sensitivity D1000 kit. Libraries were sent to the University of Michigan Sequencing Core for sequencing on the Illumina HiSeq4000 with paired-end 150 chemistry and sequenced to a depth of 50 – 70 million reads per sample. Raw reads were trimmed of low-quality bases (Phred quality score cutoff > 20) and adaptors using Trim Galore-v0.4.4 [113]. Trimmed reads were then aligned to the chicken genome, ensembl release 92, using Bismark v0.18.1 [113]. For analysis of differentially methylated cytosines (DMCs), methylkit-1.3.3 [114] was used with a read cutoff > 8x coverage and normalized, and DMCs were defined as cytosines with p-value < 0.05 and % methylation difference > 20%. Differentially methylated regions (DMRs) were identified using BSmooth v1.20.0 with a read cutoff of ≥ 2x coverage [115]. Gene ontology analysis was performed using ClueGO, a Cytoscape plug-in [116, 117], using genes with promoters overlapping with DMRs. Figures were generated using R.

## Additional data files and figures

Figure S1. Figure that demonstrate changes in the levels of 5caC in the intact RPE, t-RPE and rRPE.

Figure S2. Figure that demonstrate changes in the levels of H3K27Ac and 5mC in the intact RPE, t-RPE and rRPE.

Figure S3. depicts the pipeline used to identify DMC and DMR.

Figure S4. shows the GO networks produced identified by ClueGO.

Figure S5. demonstrate repatterning of histone and 5mC during reprogramming of RPE to NE.

Supplementary Files 1-3 contains DMRs obtained by comparing t-RPE vs RPE, rRPE vs RPE and t-rRPE vs rRPE, respectively.

Supplementary Files 4-6 contains genes with a promoter region overlapping a DMR for t-RPE vs RPE, rRPE vs RPE and t-rRPE vs rRPE, respectively.

Supplementary Files 7-9 contain GO results and networks for t-RPE vs RPE, rRPE vs RPE and t-rRPE vs rRPE, respectively.

Supplementary Table S1 contains statistical *P* values for RT-qPCR and image quantification.

Supplementary Table S2 contains the list of primers used in this study.

## Data Availability

The authors declare that all data supporting the findings of this study are available within the article and its Supplementary Information files, including Supplementary Files 1-9 or from the corresponding author upon reasonable request.

## Abbreviations

bp: Base pair
DMR: Differentially methylated region
DMC: differentially methylated cytosines
GFP: Green fluorescent protein
Histone: H3 lysine
PCR: Polymerase chain reaction
TET: Ten-eleven translocation

## Authors’ contributions

Conception and design of the study ALM and KDRT. Data acquisition, analysis, and interpretation were performed by ALM, EGE, JT, SK, LL, KW, AF, PAT and CL. Bioinformatic analysis performed by JT, LL, KW, and CL. All authors contribute to the writing of the manuscript with ALM, KDRT, EGE, and JT who were the main responsible. All authors read and approved the final manuscript.

## Acknowledgements

In loving memory to Panagiotis A. Tsonis. We acknowledge and thank the staff (Dr. Andor Kiss & Ms. Xiaoyun Deng) of the Center for Bioinformatics & Functional Genomics (CBFG) at Miami University for instrumentation and computational support, and Dr. Richard Edelmann of the Center for Advanced Microscopy and Imaging at Miami University. This work was supported by EY026816 to KDRT. We thank Leah Stetzel, Luke A. Kayafas, and Alexander McCorkle for technical contributions to this work.

