## Supplemental Figures and Tables for "DNA demethylation is a driver for chick retina regeneration"

Figure S1

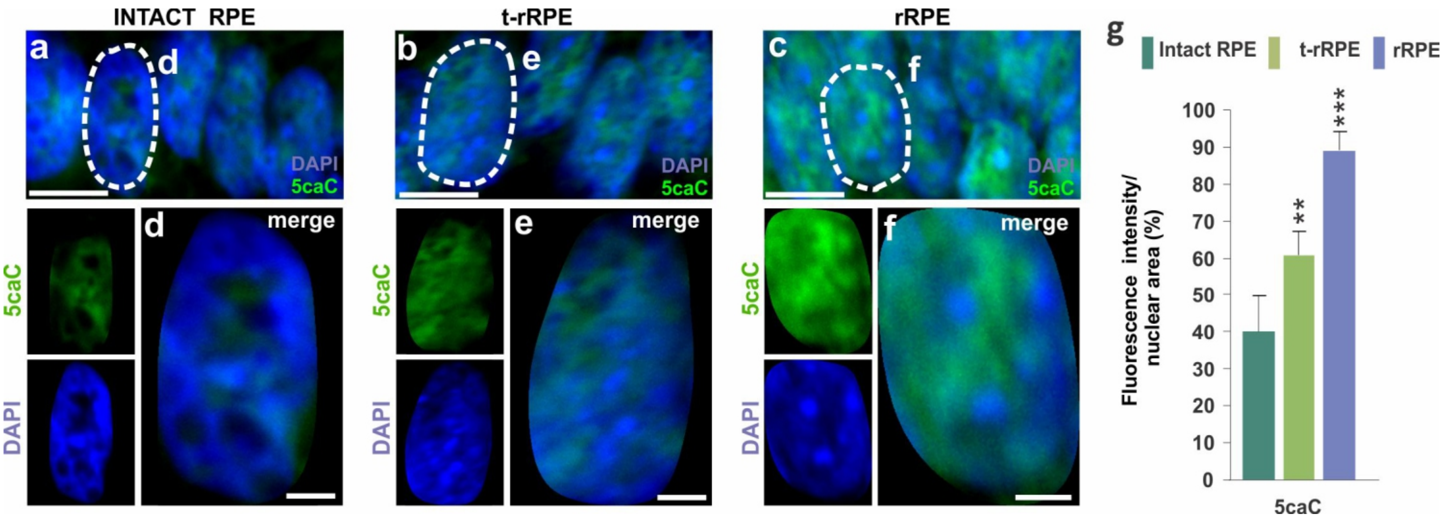

Figure S2

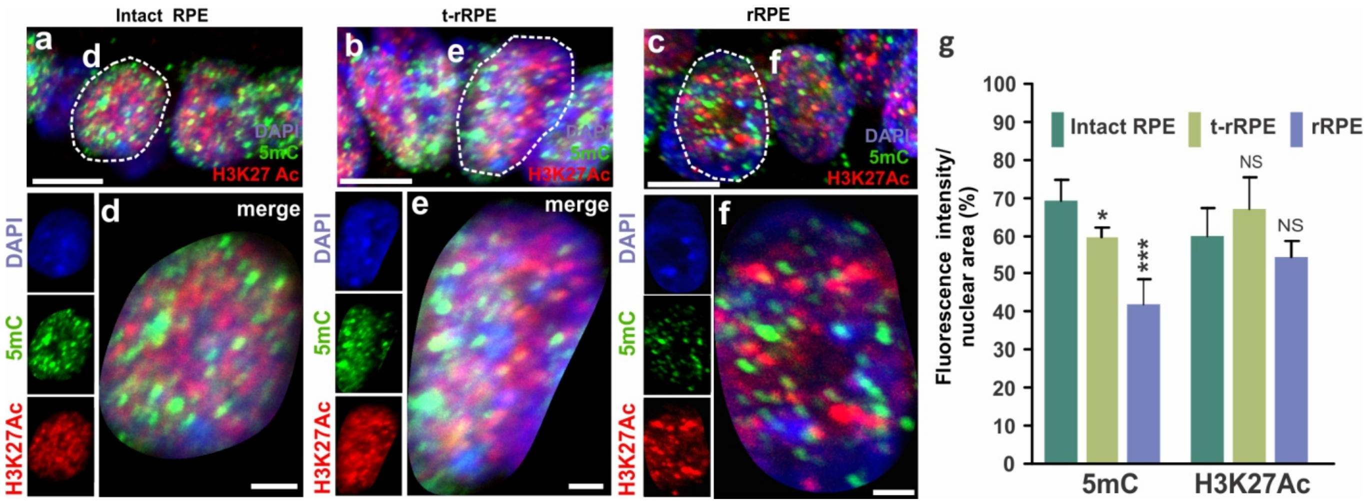

Figure S3

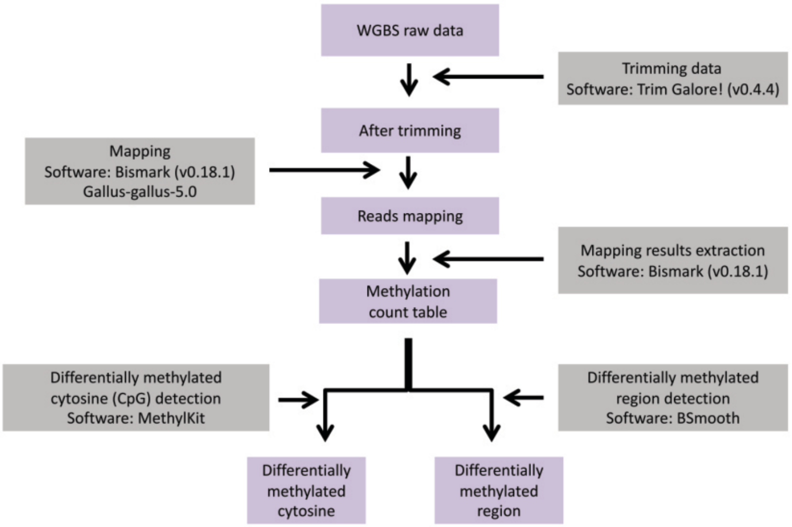

Figure S4

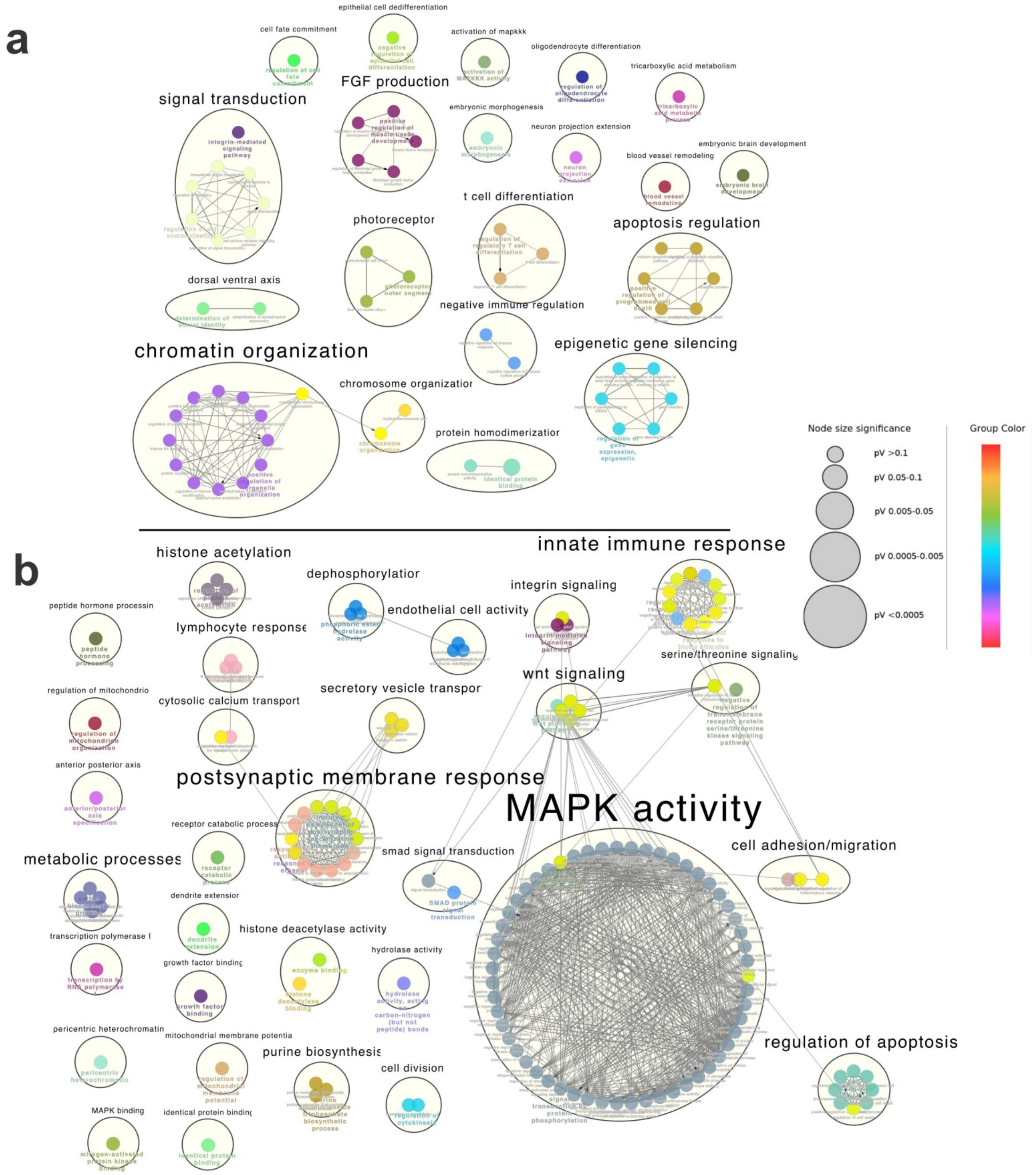
a
cell fate commitment
epithelial cell dedifferentiation
activation of mapkkk
oligodendrocyte differentiation
tricarboxylic acid metabolism
FGF production
signal transduction
embryonic morphogenesis
neuron projection extension
embryonic brain development
blood vessel remodeling
t cell differentiation
photoreceptor
apoptosis regulation
dorsal ventral axis
negative immune regulation
chromatin organization
epigenetic gene silencing
chromosome organization
protein homodimerization
Node size significance
pV >0.1
pV 0.05-0.1
pV 0.005-0.05
pV 0.0005-0.005
pV <0.0005
Group Color
b
histone acetylation
innate immune response
dephosphorylation
integrin signaling
peptide hormone processin
endothelial cell activity
lymphocyte response
serine/threonine signaling
regulation of mitochondrio
wnt signaling
cytosolic calcium transport
secretory vesicle transpor
anterior posterior axis
postsynaptic membrane response
MAPK activity
receptor catabolic process
metabolic processes
smad signal transduction
cell adhesion/migration
dendrite extension
transcription polymerase I
histone deacetylase activity
hydrolase activity
growth factor binding
pericentric heterochromatin
mitochondrial membrane potentia
purine biosynthesis
cell division
regulation of apoptosis
MAPK binding
identical protein binding

Figure S5

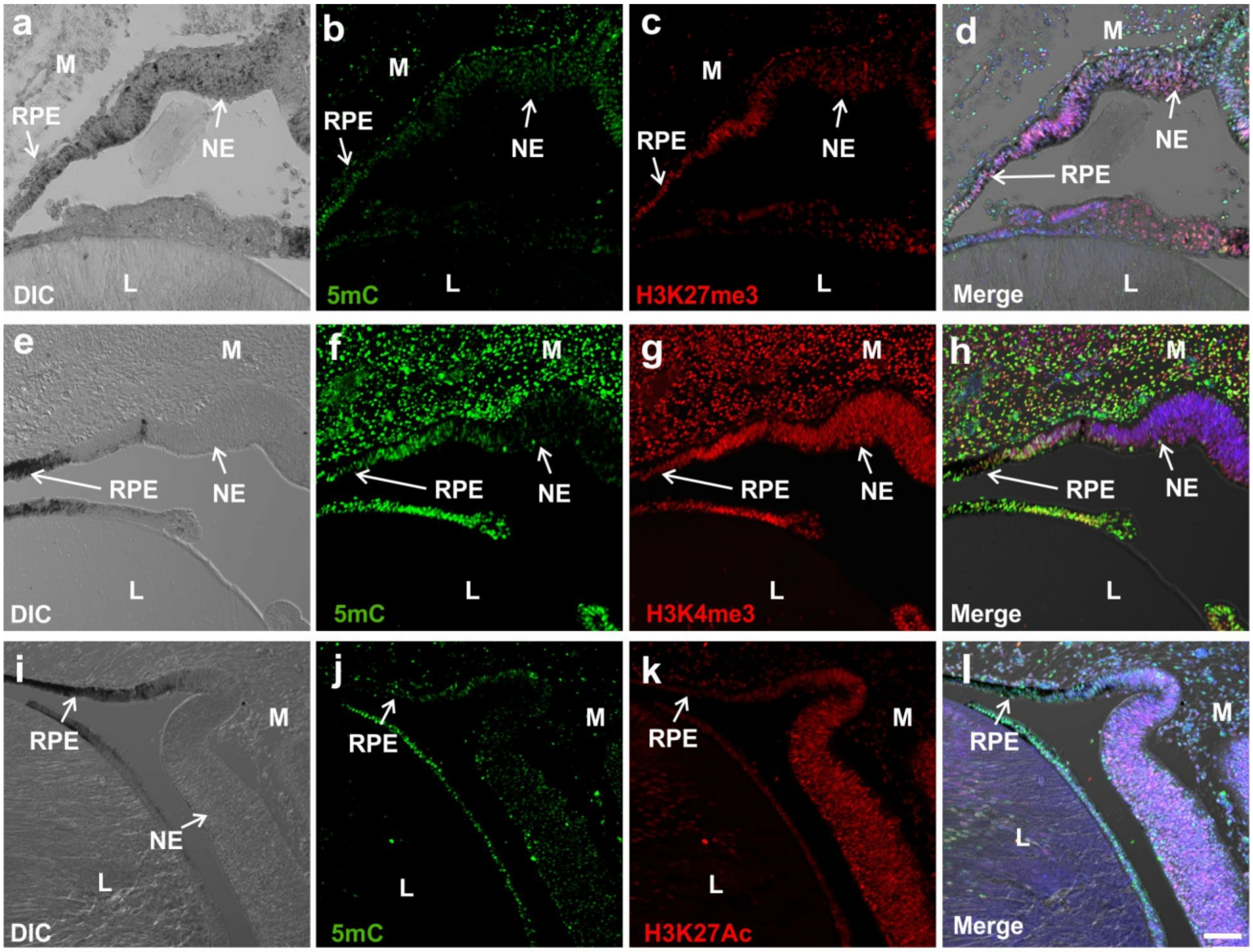

a
M
RPE
NE
DIC
L
b
M
RPE
NE
5mC
L
c
M
RPE
NE
H3K27me3
L
d
M
NE
RPE
Merge
L
e
M
RPE
NE
DIC
L
f
M
RPE
NE
5mC
L
g
M
RPE
NE
H3K4me3
L
h
M
RPE
NE
Merge
L
i
RPE
M
NE
L
DIC
j
RPE
M
L
5mC
k
RPE
M
L
H3K27Ac
l
RPE
M
L
Merge

**Supplementary Table S1**

*P* values for the percentage of fluorescence intensity per nuclear area Epigenetic marks

unpaired t-test
* = p value < 0.05
** = p value < 0.01
*** = p values < 0.001

| | |
|---|---|
| 5mC | |
| E4Dev vs Ret 6 h | 0.04152 |
| E4Dev vs Ret 6h+FGF2 | 0.00001 |
| 5hmC | |
| E4Dev vs Ret 6 h | 0.66108 |
| E4Dev vs Ret 6h+FGF2 | 0.03262 |
| 5caC | |
| E4Dev vs Ret 6 h | 0.00437 |
| E4Dev vs Ret 6h+FGF2 | 0.00002 |
| H3K27me3 | |
| E4Dev vs Ret 6 h | 0.00172 |
| E4Dev vs Ret 6h+FGF2 | 0.00371 |
| H3K4me3 | |
| E4Dev vs Ret 6 h | 0.00006 |
| E4Dev vs Ret 6h+FGF2 | 0.00015 |
| H3K27Ac | |
| E4Dev vs Ret 6 h | 0.19031 |
| E4Dev vs Ret 6h+FGF2 | 0.19309 |
| 5mC | |
| E4Ret D3+FGF2 RPE vs NE | 0.02662 |
| 5hmC | |
| E4Ret D3+FGF2 RPE vs NE | 0.00701 |
| 5caC | |
| E4Ret D3+FGF2 RPE vs NE | 0.00005 |
| H3K27me3 | |
| E4Ret D3+FGF2 RPE vs NE | 0.00043 |
| H3K4me3 | |

E4Ret D3+FGF2 RPE vs NE 0.13304

H3K27Ac
E4Ret D3+FGF2 RPE vs NE 0.00016

*P* values for relative expression RT-qPCR

unpaired t-test
* = p value < 0.05
** = p value < 0.01
*** = p values < 0.001

Dnmt1
E4Dev vs Ret 6 h 0.0001
E4 Dev vs Ret 24 h 0.0088

E4Dev vs Ret 6h+FGF2 0.0007
E4Dev vs Ret 24h+FGF2 0.0046

Dnmt3a
E4Dev vs Ret 6 h 0.0197
E4 Dev vs Ret 24 h 0.5078

E4Dev vs Ret 6h+FGF2 0.0596
E4Dev vs Ret 24h+FGF2 0.4145

Dnmt3b
E4Dev vs Ret 6 h 0.0001
E4 Dev vs Ret 24 h 0.0088

E4Dev vs Ret 6h+FGF2 0.0007
E4Dev vs Ret 24h+FGF2 0.0046

Uhrf1
E4Dev vs Ret 6 h 0.0272
E4 Dev vs Ret 24 h 0.0405

E4Dev vs Ret 6h+FGF2 0.0621
E4Dev vs Ret 24h+FGF2 0.0319

Tet1
E4Dev vs Ret 6 h 0.4527
E4 Dev vs Ret 24 h 0.8292

E4Dev vs Ret 6h+FGF2 0.4524
E4Dev vs Ret 24h+FGF2 0.0632

| | |
|---|---|
| Tet2 | |
| E4Dev vs Ret 6 h | 0.6877 |
| E4 Dev vs Ret 24 h | 0.1640 |
| E4Dev vs Ret 6h+FGF2 | 0.2820 |
| E4Dev vs Ret 24h+FGF2 | 0.3941 |
| Tet3 | |
| E4Dev vs Ret 6 h | 0.0959 |
| E4 Dev vs Ret 24 h | 0.0181 |
| E4Dev vs Ret 6h+FGF2 | 0.0121 |
| E4Dev vs Ret 24h+FGF2 | 0.0010 |
| Gadd45alpha | |
| E4Dev vs Ret 6 h | 0.0054 |
| E4 Dev vs Ret 24 h | 0.0007 |
| E4Dev vs Ret 6h+FGF2 | 0.0035 |
| E4Dev vs Ret 24h+FGF2 | 0.0048 |
| Gadd45beta | |
| E4Dev vs Ret 6 h | 0.0015 |
| E4 Dev vs Ret 24 h | 0.0004 |
| E4Dev vs Ret 6h+FGF2 | 0.0002 |
| E4Dev vs Ret 24h+FGF2 | 0.00002 |
| Gadd45gamma | |
| E4Dev vs Ret 6 h | 0.0011 |
| E4 Dev vs Ret 24 h | 0.0003 |
| E4Dev vs Ret 6h+FGF2 | 0.00005 |
| E4Dev vs Ret 24h+FGF2 | 0.00015 |
| tdg | |
| E4Dev vs Ret 6 h | 0.0003 |
| E4 Dev vs Ret 24 h | 0.0036 |
| E4Dev vs Ret 6h+FGF2 | 0.0005 |
| E4Dev vs Ret 24h+FGF2 | 0.00002 |
| Prdm1 | |
| E4Dev vs Ret 6 h | 0.0015 |
| E4 Dev vs Ret 24 h | 0.0041 |

| | |
|---|---|
| E4Dev vs Ret 6h+FGF2 | 0.0015 |
| E4Dev vs Ret 24h+FGF2 | 0.0001 |

Suz12

| | |
|---|---|
| E4Dev vs Ret 6 h | 0.7675 |
| E4 Dev vs Ret 24 h | 0.8379 |
| E4Dev vs Ret 6h+FGF2 | 0.6811 |
| E4Dev vs Ret 24h+FGF2 | 0.6056 |

eed

| | |
|---|---|
| E4Dev vs Ret 6 h | 0.3283 |
| E4 Dev vs Ret 24 h | 0.7555 |
| E4Dev vs Ret 6h+FGF2 | 0.3489 |
| E4Dev vs Ret 24h+FGF2 | 0.0139 |

ezh2

| | |
|---|---|
| E4Dev vs Ret 6 h | 0.8094 |
| E4 Dev vs Ret 24 h | 0.9369 |
| E4Dev vs Ret 6h+FGF2 | 0.3914 |
| E4Dev vs Ret 24h+FGF2 | 0.3068 |

Jmjd1

| | |
|---|---|
| E4Dev vs Ret 6 h | 0.1381 |
| E4 Dev vs Ret 24 h | 0.0034 |
| E4Dev vs Ret 6h+FGF2 | 0.0461 |
| E4Dev vs Ret 24h+FGF2 | 0.0236 |

Jhdm1d

| | |
|---|---|
| E4Dev vs Ret 6 h | 0.0031 |
| E4 Dev vs Ret 24 h | 0.0647 |
| E4Dev vs Ret 6h+FGF2 | 0.2176 |
| E4Dev vs Ret 24h+FGF2 | 0.6863 |

Jmjd1c

| | |
|---|---|
| E4Dev vs Ret 6 h | 0.7830 |
| E4 Dev vs Ret 24 h | 0.6175 |

| | |
|---|---|
| E4Dev vs Ret 6h+FGF2 | 0.9779 |
| E4Dev vs Ret 24h+FGF2 | 0.4105 |

Jmjd4

| | |
|---|---|
| E4Dev vs Ret 6 h | 0.4089 |
| E4 Dev vs Ret 24 h | 0.6564 |
| E4Dev vs Ret 6h+FGF2 | 0.9970 |
| E4Dev vs Ret 24h+FGF2 | 0.6020 |

Jmjd5

| | |
|---|---|
| E4Dev vs Ret 6 h | 0.1846 |
| E4 Dev vs Ret 24 h | 0.0161 |
| E4Dev vs Ret 6h+FGF2 | 0.00001 |
| E4Dev vs Ret 24h+FGF2 | 0.4976 |

Kdm5b

| | |
|---|---|
| E4Dev vs Ret 6 h | 0.3387 |
| E4 Dev vs Ret 24 h | 0.0063 |
| E4Dev vs Ret 6h+FGF2 | 0.8396 |
| E4Dev vs Ret 24h+FGF2 | 0.2308 |

Utx

| | |
|---|---|
| E4Dev vs Ret 6 h | 0.3032 |
| E4 Dev vs Ret 24 h | 0.3208 |
| E4Dev vs Ret 6h+FGF2 | 0.2514 |
| E4Dev vs Ret 24h+FGF2 | 0.1817 |

**Supplementary Table S2.**-Primer sequences for RT-qPCR

| **Gene name** | **ENSEMBL ID or NCBI Reference Sequence** | **Forward primer (5'-3')** | **Reverse primer (5'-3')** |
|---|---|---|---|
| *dnmt1* | ENSGALT00000051245.2 | TGTCCATCTTCGACGCCAAC | CATAGATGGGCTTCACGGCA |
| *dnmt3a* | ENSGALT00000063127.3 | GGGACGGCAAGTTCTCAGT | ATGGGCTGCTTGTTGTAGGT |
| *dnmt3b* | ENSGALT00000099598.1 | GCCACGTTCAATAAGCTGGT | AGGGGCAGATGTGAATGTCT |
| *uhrf1* | ENSGALT00000018849.6 | GAAAGCCATGGTGCAGTATG | GTGTTGTTTCAGAAGCCAAGC |
| *tet1* | ENSGALT00000060823.2 | GCTATGCTTGGTACGTCAGC | TCCCACGCCAGTATGAGAAT |
| *tet2* | ENSGALT00000108375.1 | CCTGCCAAAACCCAGTATGA | AGGTGGGTATAGAAAGGACCT |
| *tet3* | XM_015297468.2 | GGAAGGAGGGGAAGAGCTC | GCACAGCAGCTTCTCCTC |
| *gadd45α* | ENSGALT00000046015.3 | GTCTACGAGGCGGCCAAG | TGAAGTGGATTTGCAGAGCC |
| *gadd45β* | XM_015299957.2 | CTGAGCCCCGTGATCTGC | GCTCTCGGCGCAGTAATTC |
| *gadd45γ* | ENSGALT00000045592.3 | GTCGGCCAAGCTTATGAACG | TGAAGTGGATCTGCAGGGC |
| *tdg* | ENSGALT00000052099.2 | TGGCTGTCAAGGAAGAAAAGT | TTTCATATGGCGCACGATCG |
| *prdm14* | XM_025147991.1 | TCCACGAGAAGCACAGACC | GAGTGAACCCGCATGTGTTT |
| *suz12* | XM_004946169.3 | TTTGCCACAAGAAACGAAAG | TGCCTGGTTTGATTTGACTG |
| *eed* | ENSGALT00000054939.3 | CTAATGCACCGGGAAGAAAA | GGACTCCAAACAAAGGCTGA |
| *ezh2* | ENSGALT00000072028.2 | GGAGAAAACAATGATAAAGAGGAA | TGTTTGACACCTAGAATTTGCTT |
| *jmjd1a* (*kdm3a*) | ENSGALT00000037139.4 | TGATGCCAAACACACGAAGT | GCAGGAGTACAACCCAGCA |
| *jhdm1d* (*kdm7a*) | ENSGALT00000060010.3 | ATCCTGACCGACCAAAAGTG | AATTTTCCACCCATGAAAGC |
| *jmjd1c* | ENSGALT00000004648.6 | ATGATCTCTTCACCCGCACT | CAACAAAGAGTGAACAACGTCA |
| *jmjd4* | ENSGALT00000008701.7 | TCCGACTGGCTGAATGAGTA | ACGTCAGCATGGAATGGAGT |
| *jmjd5* (*kdm8*) | ENSGALT00000010182.6 | ACCGCTACCTCATTCCACAG | ACTTGACGCACGTAGTCCAC |
| *jarid1b* (*kdm5b*) | ENSGALT00000056356.2 | TTTAATTCCACCCCTTCACG | ACGCTTCCTGAGGCTTATTG |
| *utx* (*kdm6a*) | ENSGALT00000026143.5 | TTTTGTCGAGCCAAGGAAAT | AGGGGTTGCAGTCAATCAAA |
| gapdh | | | |
